# Cryo-EM structures of calcium homeostasis modulator channels in diverse oligomeric assemblies

**DOI:** 10.1101/2020.01.31.928093

**Authors:** Kanae Demura, Tsukasa Kusakizako, Wataru Shihoya, Masahiro Hiraizumi, Kengo Nomura, Hiroto Shimada, Keitaro Yamashita, Tomohiro Nishizawa, Akiyuki Taruno, Osamu Nureki

## Abstract

Calcium homeostasis modulator (CALHM) family proteins are Ca^2+^-regulated ATP-release channels involved in neural functions including neurotransmission in gustation. Here we present the cryo-EM structures of killifish CALHM1, human CALHM2, and *C. elegans* CLHM-1 at resolutions of 2.66, 3.51, and 3.60 Å, respectively. The CALHM1 octamer structure reveals that the N-terminal helix forms the constriction site at the channel pore in the open state, and modulates the ATP conductance. The CALHM2 undecamer and CLHM-1 nonomer structures show the different oligomeric stoichiometries among CALHM homologs. We further report the cryo-EM structures of the chimeric construct, revealing that the inter-subunit interactions at the transmembrane domain define the oligomeric stoichiometry. These findings advance our understanding of the ATP conduction and oligomerization mechanisms of CALHM channels.

**One Sentence Summary:** Cryo-EM structures reveal the ATP conduction and oligomeric assembly mechanisms of CALHM channels.

## Introduction

Adenosine triphosphate (ATP) as an extracellular ligand plays essential roles in various cellular functions, including neurotransmission (*1*), (*2*). Although anionic ATP molecules cannot diffuse across the plasma membrane, ATP release from the cytosol is mediated by ATP-permeable channels, such as connexin hemichannels, pannexin 1 (PANX1), volume-regulated anion channels (VRACs), and calcium homeostasis modulator 1 (CALHM1) (*3*). CALHM1 was originally identified as a genetic risk factor for late-onset Alzheimer’s disease (*4*), and recently identified as an essential component of the neurotransmitter-release channel in taste bud cells that mediates the voltage-dependent release of ATP to afferent gustatory neurons, whereby mediating the perception of sweet, bitter, and umami tastes (*5*), (*6*), (*7*). CALHMs are regulated by the membrane voltage and extracellular Ca^2+^. Strong de-polarization and extracellular Ca^2+^ concentration ([Ca^2+^]_o_) reduction increase the open probability of CALHM channels (*8*), (*9*). Six CALHM1 homologs were found in humans (*10*). CALHM genes are present throughout vertebrates, but they have no sequence homology to other known genes. CALHM2 modulates neural activity in the central nervous system (*11*). CALHM3 interacts with CALHM1, forming a CALHM1/CALHM3 heteromeric channel that confers fast activation (*12*), (*13*). Outside of vertebrates, *Caenorhabditis elegans* (*C. elegans*) possesses single CALHM ortholog, termed CLHM-1. The *C. elegans* CLHM-1 is expressed in sensory neurons and muscles, and functions as an ion channel (*14*), (*15*).

Many ATP-permeable channels have been structurally characterized (*16*), (*17*), (*18*), (*19*), but the molecular mechanism of CALHM channels has remained unclear. Here we report the cryo-EM structures of three CALHM homologs (CALHM1, CALHM2, and *C. elegans* CLHM-1) and their chimeric construct in unexpectedly different oligomeric states, revealing the oligomerization mechanism of CALHM channels. Furthermore, the present cryo-EM structure of CALHM1 is the first to visualize the N-terminal helix, which forms the constriction gate at the channel pore in the open state, and the bioluminescence assay revealed that it modulates the ATP conductance. These findings provide insights into the ATP conduction and oligomerization mechanisms of CALHM channels.

## Structure determination of CALHM1

For the structural analysis, we first screened the CALHM1 proteins from various vertebrates by fluorescence detection gel filtration chromatography (FSEC) (*20*), and identified the killifish *Oryzias latipes* CALHM1 (OlCALHM1) as a promising candidate (Fig. S1A). OlCALHM1 and HsCALHM1 (’Hs’ refers to *Homo sapiens*) share 62% sequence identity and 78% sequence similarity within the transmembrane region. Next, we analyzed the function of OlCALHM1. We transfected the DNA encoding the full-length OlCALHM1 into HeLa cells, and measured the extracellular ATP concentration by the luciferin–luciferase reaction (*5*) (Fig. S2A–C). Lowering [Ca^2+^]_o_ induced a larger increase in the extracellular ATP concentration than in mock-transfected cells (Fig. S2A), and ruthenium red (RUR), a known blocker of human and mouse CALHM1 channels, abolished this ATP release (Fig. S2B), indicating that OlCALHM1 forms an ATP-release channel activated by low [Ca^2+^]_o_, as also reported for HsCALHM1 (*5*).

For the cryo-EM analysis, we expressed the CALHM protein in HEK293S GnT1^-^ cells. To facilitate the expression and purification of OlCALHM1, we truncated the flexible C-termini starting from Trp296 (OlCALHM1_EM_). OlCALHM1_EM_ retains the ATP-release activity elicited by low [Ca^2+^]_o_ when expressed in HeLa cells (Fig. S2C). Moreover, we confirmed that the purified OlCALHM1_EM_ forms an ATP-permeable channel by a liposome assay (Fig. S2D). The OlCALHM1_EM_ protein was purified in GDN micelles without Ca^2+^ and vitrified on a grid. The samples were imaged using a Titan Krios electron microscope equipped with a K3 camera. The 2D class average showed obvious C8 symmetry (Fig. S3A, B). The 3D map was reconstructed from 102,109 particles with the imposed C8 symmetry, at the 2.66 Å resolution based on the Fourier shell correlation (FSC) gold standard criteria (Fig. 1A), which was sufficient for *de novo* model building (Fig. S3C–E, Fig. S4A and Table S1). The processing summary is provided in Fig. S3B. The N-terminal residues Met1 to Met6 and the C-termini after Gln263 are disordered, consistent with the secondary structure prediction.

**Fig. 1:**
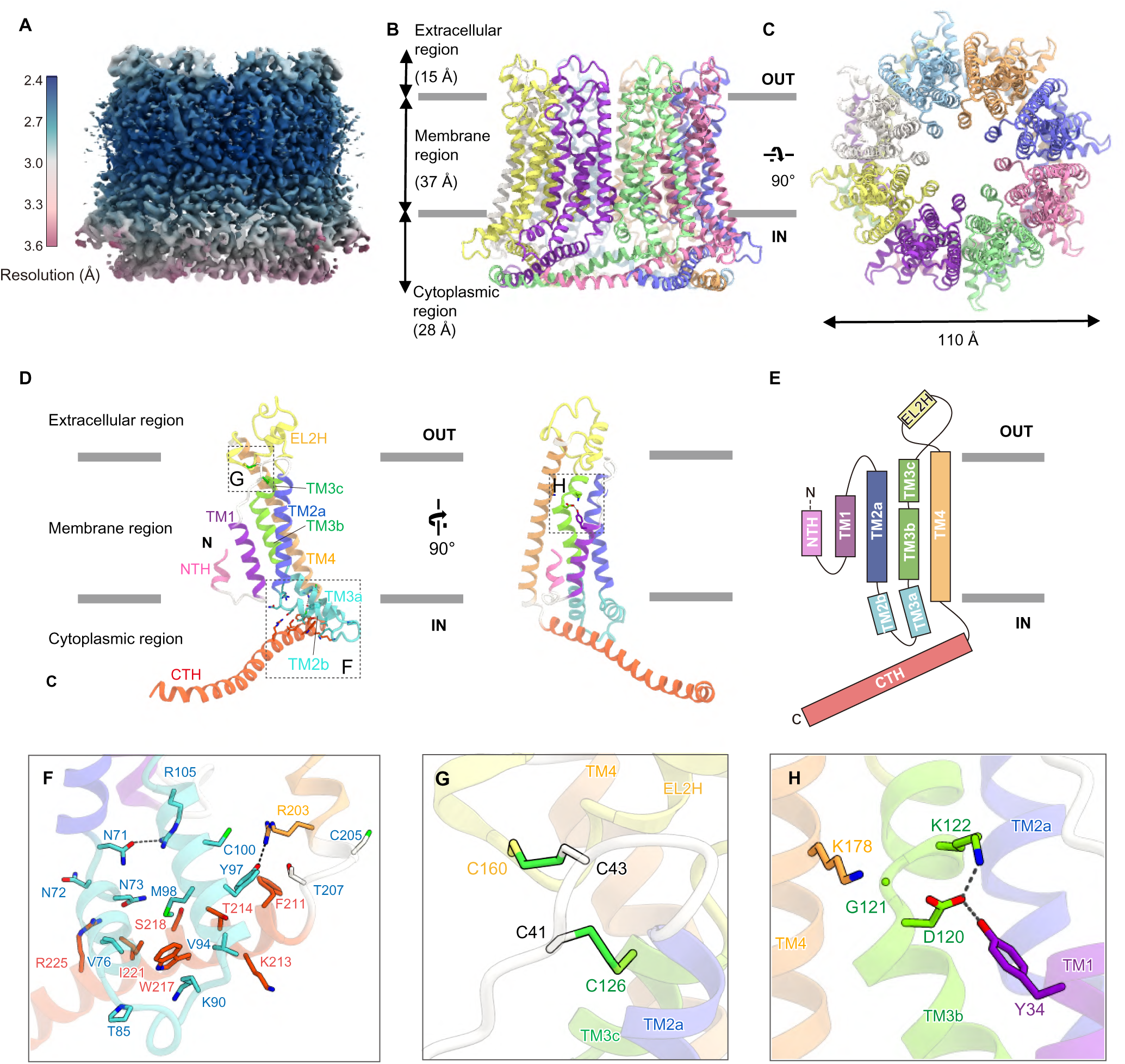
Overall structure of OlCALHM1. (**A**) Local resolution of the OlCALHM1_EM_ structure, estimated by RELION. (**B** and **C**) Overall structure of the OlCALHM1_EM_ octamer, viewed parallel to the membrane (B) and from the extracellular side (C). (**D**) The subunit structure of OlCALHM1_EM_, viewed parallel to the membrane. Each region is colored as follows: NTH, pink; TM1, purple; EL1, white; TM2a, light blue; TM2b, IL1, and TM3a, cyan; TM3b and 3c, light green; EL2, yellow; TM4, orange; CTH, red. The N- and C-termini are indicated by ‘N’ and ‘C’, respectively. (**E**) Schematic representation of the OlCALHM1 membrane topology, colored as in Fig. 1D. (**F**–**H**) Close-up views of the intracellular side (F), the two disulphide bonds in the extracellular region (G), and the interactions between TM1 and the other TMs around Asp120 (H).

## Structure of OlCALHM1

OlCALHM1_EM_ forms an octamer with a length of 80 Å and a width of 110 Å (Fig. 1B, C). At the center of the octamer structure, a channel pore is formed along an axis perpendicular to the membrane. The extracellular region projects approximately 15 Å above the plasma membrane. The transmembrane region is 37 Å thick, and the intracellular region protrudes about 28 Å from the plasma membrane. The subunit consists of extracellular loops 1 and 2 (EL1: Phe37–Tyr47 and EL2: Phe128–Asp166), an intracellular loop (IL: Arg83–Ala91), an N-terminal helix (NTH: Met7–Gln15), a transmembrane domain (TMD: Gly25–Ser35, Tyr48–Leu70, Ile108–Ala127, and Asn167–Cys205), and a C-terminal helix (CTH: Phe206–Arg260), with the C-terminus on the cytoplasmic side (Fig. 1D, E). The TMD consists of four transmembrane helices (TM1–4), which are aligned counterclockwise (Figs. 1D, E and S4B) as viewed from the extracellular side. This TM arrangement is quite different from those in the connexin (*16*), innexin (*17*), and LRRC8 (*18*), (*19*) structures (Fig. S4B). TM1 faces the channel pore, while TM2 and TM4 face the phospholipid membrane. A clear density shows the N-terminal helix (NTH) preceding TM1 and forming the channel pore, together with TM1 (Fig. S4A). TM3 is located between TMs 1, 2, and 4, connecting these structural elements. TM4 is tilted by about 30° from the channel pore axis (perpendicular to the membrane plane), and extends from the membrane lipid. Notably, TM2 is kinked at Asn71 (TM2a and 2b), and TM3 is kinked at Leu107 and Gly121 (TM3a-c). Thereby, TM2b and TM3a protrude into the intracellular region, and are stabilized by the hydrogen bonding interactions of Asn71–Arg105 and Tyr97–Arg203. On the intracellular side, CTH adopts a long helix and extends toward the adjacent subunit. The protruded TM2b and TM3a form extensive van der Waals interactions with the CTH (Fig. 1F). On the extracellular side, EL2 between TM3 and TM4 forms a short helix (EL2H) and wraps around the short EL1 between TM1 and TM2, covering the extracellular side of the transmembrane region. Between EL1 and EL2, two disulfide bonds are formed at the Cys41–Cys126 and Cys43– Cys160 pairs, which stabilize the EL structure (Fig. 1G). The residues constituting the transmembrane domain and the CTH, and the two cysteines in the extracellular loops are highly conserved in the CALHM1 orthologs, including HsCALHM1 (Fig. S1A–C).

A previous study reported that two cysteines in mouse CALHM1 are palmitoylated and associated with the post-translational regulation of the channel gating and localization (*21*). These two cysteines are completely conserved in the CALHM1 orthologs (Fig. S1A). In the OlCALHM1_EM_ structure, the corresponding cysteines Cys100 and Cys205 are located at TM3a and the intracellular end of TM4, respectively (Fig. 1F). Although we could not find the density of the palmitoylation in our EM map, these residues face the lipid bilayer and are likely to be palmitoylated.

The P86L polymorphism in HsCALHM1 is associated with Alzheimer’s disease (*4*). In the OlCALHM1_EM_ structure, the corresponding residue Thr85 is located on the intracellular loop between TM2b and TM3a. Therefore, Thr85 is not oriented toward the channel pore and not directly associated with the TMD-CTH interface. Given that the palmitoylation site is also located on TM2b (Fig. 1F), the protruded TM2b and TM3a probably have an important role in channel function.

In HsCALHM1, the negative charge of Asp121 reportedly plays a critical role in ion permeation property (*8*). This Asp residue is highly conserved in the CALHM1 orthologs, and the corresponding Asp120 in OlCALHM1 forms a salt bridge with Lys122 in TM3 (Fig. 1H), stabilizing its bent conformation at Gly121. Moreover, Asp120 forms a hydrogen bond with Tyr34 in TM1 and an electrostatic interaction with Lys178 in TM4. These residues are also conserved in HsCALHM1. Overall, Asp120 contributes to the structural organization of the TM helices inside the channel.

## Subunit interface in OlCALHM1

The subunit interface is formed at the transmembrane and intracellular regions (Fig. 2A). Unlike connexin (*16*), innexin (*17*), and LRRC8 (*18*), (*19*), the extracellular region is not involved in the subunit interface. At the transmembrane region (Fig. 2B), TM2 forms extensive hydrophobic interactions with TM4’ (apostrophe indicates the adjacent subunit). At the extracellular and intracellular ends of the interface, Tyr51 and Asn72 in TM2 form hydrogen bonds with Gln182 and Arg200 in TM4’, respectively (Fig. 2B). EL1 also forms hydrogen-bonding interactions with TM4’ (Fig. 2C). Moreover, the NTH regions interact with each other between adjacent subunits: Gln12 in the NTH forms a hydrogen bonding interaction with Ser13 in the NTH’ (Fig. 2C). At the intracellular region, the long CTH extends to the adjacent subunit and forms extensive interactions with it (Fig. 2D). Furthermore, Glu254 forms a salt bridge with Lys213 in the CTH between the two adjacent subunits (Fig. 2D).

**Fig. 2:**
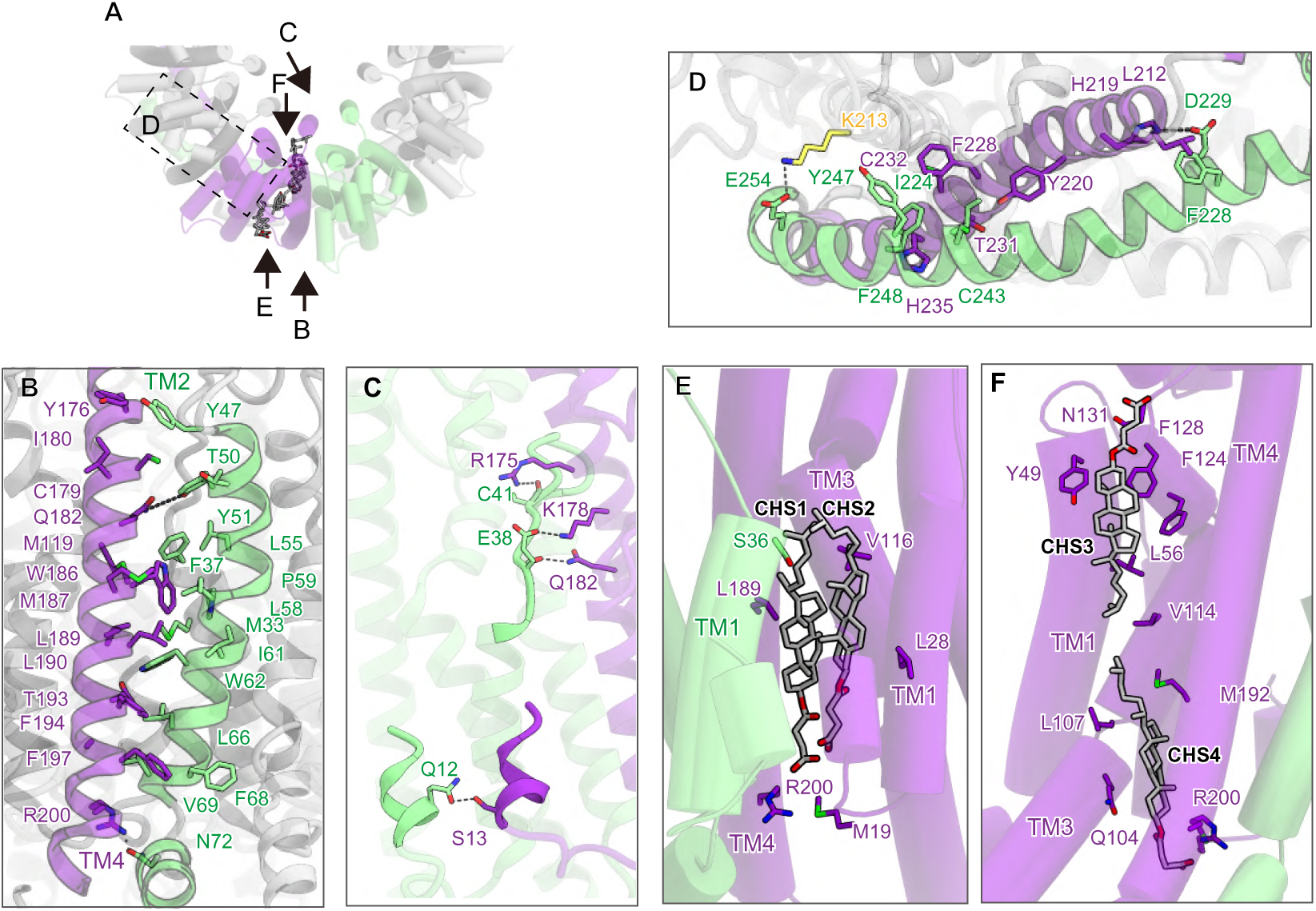
Subunit interfaces between two adjacent subunits. (**A**) Overview between two adjacent subunits, viewed from the extracellular side. (**B**–**D**) Close-up views between two adjacent subunits at the transmembrane region (B and C) and intracellular region (D). (**E** and **F**) Close-up views of CHS1, 2 (E) and CHS3, 4 (F). CHS molecules are indicated by grey stick models.

In addition, there are extra densities at the subunit interface (Figs. 2E, F and S4A). Based on their characteristic shape, we assigned them to the cholesteryl hemisuccinate (CHS) molecules used for solubilization (CHS1, 2) (Fig. 2E). These CHS molecules are surrounded by TM1 (Ser36) and TM1’ (Met28 and Leu19), TM3 (Val116), and TM4 (Arg200 and Leu189), and bridge the gap between the subunits behind the NTH, stabilizing the subunit interface and pore architecture. A lipid-mediated subunit interface has not been observed in connexin26, LRRC8, or other gap junction channels, and is a unique structural feature of OlCALHM1. Moreover, two CHS molecules are observed on the membrane side (CHS3, 4) (Fig. 2F). These CHS molecules form extensive hydrophobic interactions with the channel and fill the cleft between TMs1, 3, and 4. The residues involved in the interaction with CHS are highly conserved in the CALHM1 orthologs (Fig. S1), suggesting that lipids play an important role in the structural stabilization of the CALHM1 channels.

## Pore architecture of OlCALHM1

OlCALHM1_EM_ has a channel pore along a central axis perpendicular to the membrane. In the structures of gap junction channels (connexin and innexin) (*16*), (*17*), the NTH and extracellular loop form two constriction sites in the pore. In the OlCALHM1_EM_ structure, only the NTH forms the single constriction site in the pore (Fig. 3A). At the constriction site, the neutral residues Gln9, Gln12, and Ser13 face the pore, with the narrowest diameter of 15.7 Å at Gln9 (Fig. 3B). Previous optical and electrophysiological analyses of the permeation of fluorescence dyes and ions with different sizes suggested that HsCALHM1 has a large pore greater than 14 Å (*10*), which is consistent with the diameter at the narrowest neck of OlCALHM1_EM_, allowing the permeation of all hydrated ions and ATP (∼7 Å). HsCALHM1 is both cation and anion permeable at P_Na_: P_K_: P_Ca_: P_Cl_ = 1: 1.14: 13.8: 0.52 (*8*), (*10*). The pore-facing residues in the NTH are neutral (Gln9, Gln12, and Ser13), accounting for the weak charge selectivity. Taken together, the pore architecture in the OLCALHM1_EM_ structure is in excellent agreement with the previous functional analysis of HsCALHM1.

**Fig. 3:**
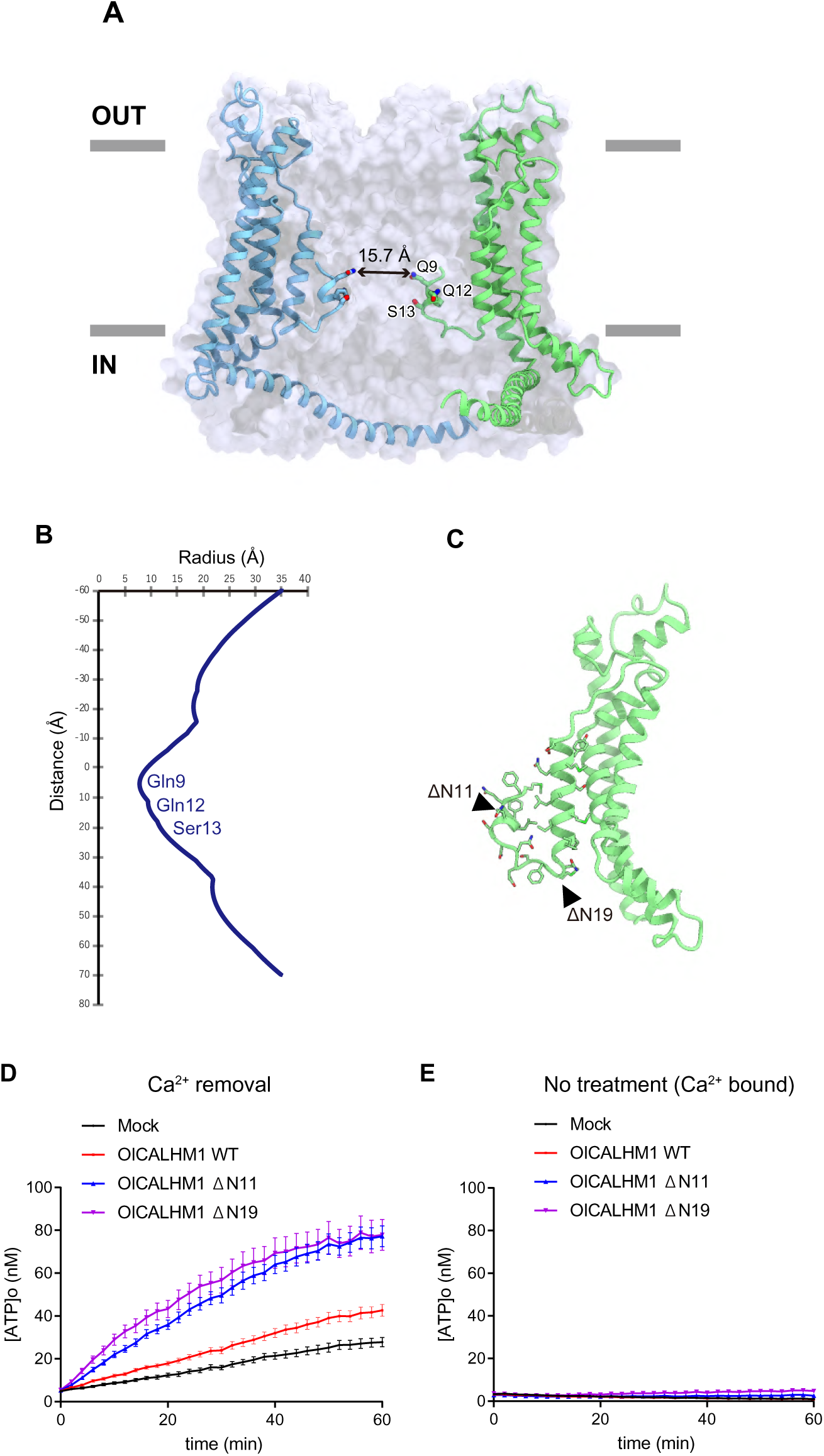
Pore architecture and ATP conduction of OlCALHM1. (**A**) Cross-sections of the surface representation for the OlCALHM1_EM_ subunits, showing the ion channel pore. (**B**) The pore radius for the OlCALHM1_EM_ structure along the pore center axis. The pore size was calculated with the program HOLE (*23*). (**C**) Design of the ΔNTH constructs. The residues in the NTH and the loop between NTH and TM1 are shown as stick models. (**D** and **E**) Time courses of extracellular ATP levels due to release from HeLa cells transfected with the empty vector (mock), WT OlCALHM1, or NTH-truncated (ΔN11 or ΔN19) OlCALHM1 following exposure to essentially zero (17 nM) (D) or normal (1.9 mM) [Ca^2+^]_o_ (E). Data are displayed as means ± S.E.M. n = 15 (Mock), n = 16 (WT, ΔN11, and ΔN19) in D. n = 16 in each group in E.

To elucidate the functional role of the NTH, we truncated the NTH of OlCALHM1 (ΔN11 and ΔN19) (Fig. 3C) and measured the ATP release activity by a cell-based assay. In the Ca^2+^-free open state, the ATP release activities of the NTH-truncated mutant channels were markedly enhanced as compared to the wild-type, indicating that the NTH importantly functions in the ATP release (Fig. 3D). However, in the Ca^2+^-bound closed state, the NTH-truncated mutants showed little ATP release activities (Fig. 3E). These observations indicate that the NTH observed in our CALHM1 structure is indeed involved in the channel conduction of ATP; however, the other structural components are associated with the Ca^2+^-dependent channel closing.

## Structures of HsCALHM2 and CeCLHM-1 and structural differences among CALHM homologs

Next, we investigated the functional and structural diversity of the CALHM family. Although various members of the CALHM family have been identified (*11*), (*14*), (*15*), their ATP conductance activities remain elusive. Thus, we measured the ATP release activities of HsCALHM2 and CeCLHM-1 by a cell-based assay. HsCALHM2- and CeCLHM-1-transfected HeLa cells exhibited greater RuR-sensitive increases in the extracellular ATP concentration in response to lowering [Ca^2+^]_o_ compared to mock-transfected cells, indicating that HsCALHM2 and CeCLHM-1 form ATP-release channels activated by low [Ca^2+^]_o_, as in the CALHM1 channels (Fig. S2A, B). To further understand the ATP conduction mechanisms of the CALHM family members, we performed the cryo-EM analysis of HsCALHM2 and CeCLHM-1. HsCALHM2 and CeCLHM-1 were purified without Ca^2+^ and exchanged into the amphipol PMAL-C8. The cryo-EM images were acquired on a Titan Krios electron microscope equipped with a Falcon 3 direct electron detector. The 2D class averages of HsCALHM2 and CeCLHM-1 showed the top views of 11-mer and 9-mer, respectively. We finally reconstructed the EM density maps of HsCALHM2 and CeCLHM-1 at resolutions of 3.51 Å and 3.6 Å, respectively, according to the gold-standard Fourier shell correlation (FSC) = 0.143 criteria (Figs. S5, S6 and Table S1).

Unlike the OlCALHM1 8-mer structure, HsCALHM2 and CeCLHM-1 form the 11-mer (approximate dimensions of 93 Å in height × 143 Å in intracellular width) and the 9-mer (approximate dimensions of 93 Å in height ×126 Å in intracellular width), respectively (Fig. 4A). The 11-mer structure of HsCALHM2 is similar to that of the recently reported structure in the open state (*22*). Each subunit of the three homologs, OlCALHM1, HsCALHM2, and CeCLHM-1, adopts almost similar architectures composed of the transmembrane domain (TMD) and the C-terminal domain (CTD) (Fig. 4B, C). At the TMD, both HsCALHM2 and CeCLHM-1 have the 4-TM helix topology, in which the NTH and TM1 of HsCALHM2 and CeCLHM-1 are also expected to face the pore, similar to OlCALHM1. However, the NTH and TM1 are not visualized in the density maps of HsCALHM2 and CeCLHM-1, indicating the conformational flexibility of these regions. At the extracellular regions, two disulfide bonds between extracellular loops 1 (EL1) and 2 (EL2) are conserved among the three CALHM homologs (Fig. S7). EL2, between TM3 and TM4, forms short helices that are arranged differently in the three structures. At the CTD, the three CALHM homologs adopt distinct architectures (Fig. 4B, C). In contrast to OlCALHM1, HsCALHM2 and CeCLHM-1 have a CTH and additional helices, and their arrangements are different between the two homologs (Fig. 4B, C): The CTD of HsCALHM2 consists of a CTH (Arg215–Phe250), which turns toward the pore at Phe251, and three short helices. The CTD of CeCLHM-1 consists of a CTH (Leu220– Arg254), which turns toward the membrane at Phe257, and at least one short helix. In addition to these structural observations, the low sequence identities within the CTDs among the three homologs (OlCALHM1 vs. HsCALHM2: 23%, OlCALHM1 vs. CeCLHM-1: 32%) suggest the structural divergence of the CTD among the CALHM family members.

**Fig. 4:**
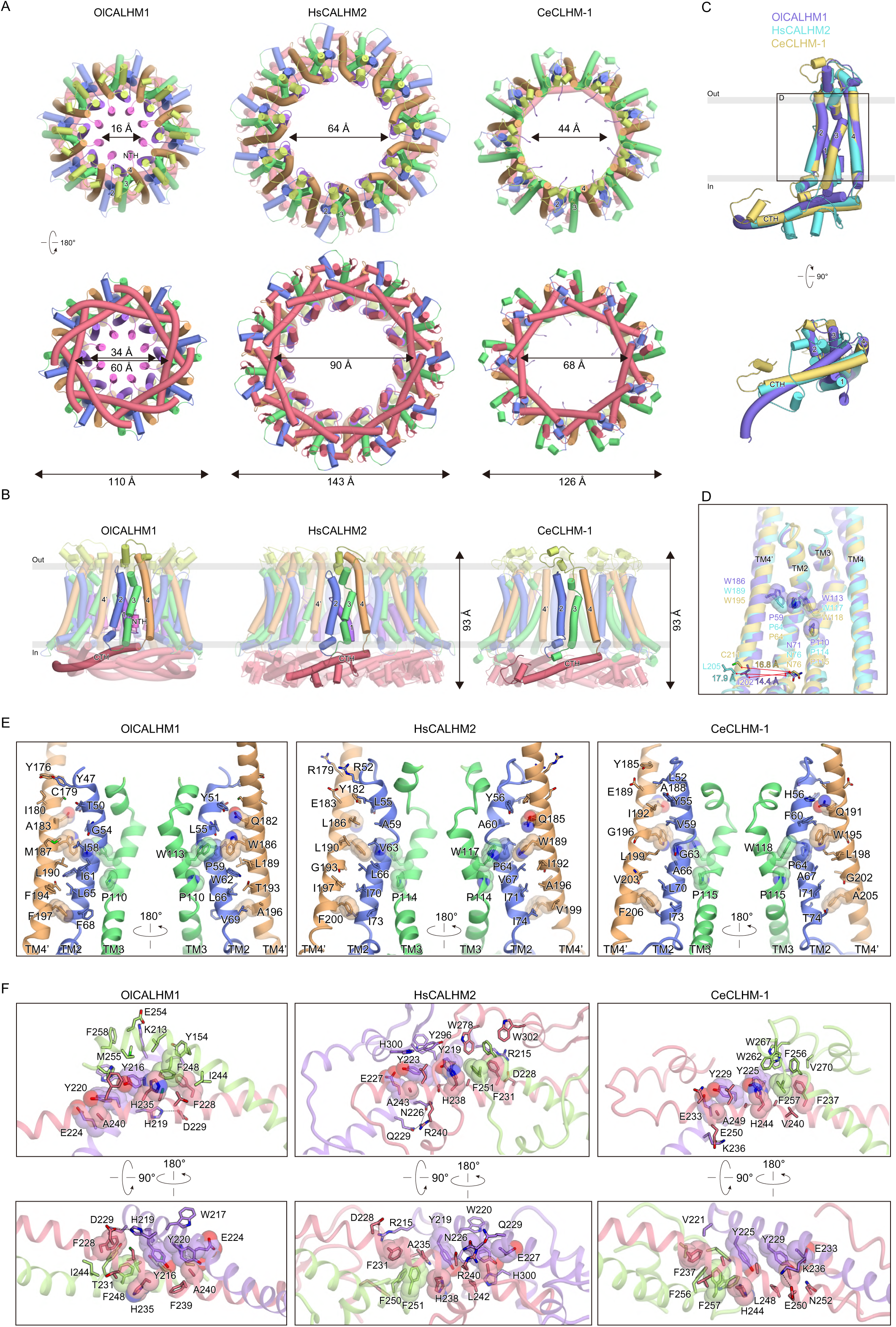
Structural differences among OlCALHM1, HsCALHM2, and CeCLHM-1. (**A**) Structural comparisons among OlCALH1, HsCALHM2, and CeCLHM1. Overall structures are viewed from the extracellular side (top) and the intracellular side (bottom), colored as in Fig. 1D. (**B**) Subunit structures of the three CALHM channels, colored as in Fig. 1D. Adjacent subunits are shown in semi-transparent colors. (**C**) Superimposition of the subunit structures of the three CALHM channels, viewed from the membrane side (left) and the intracellular side (right). OlCALHM1, HsCALHM2, and CeCLHM-1 are colored purple, cyan, and yellow, respectively. (**D**) Close-up view of the superimposition of the subunit structures of the three CALHM channels, colored as in Fig. 4C. Red arrows indicate the different orientations of TM4’ in the adjacent subunits among the three CALHM structures. (**E**) Subunit interfaces at the TMD of the three CALHM channels. TM2, TM3, and TM4 are colored blue, green, and orange, respectively. The conserved residues among the CALHM channels are shown in stick models and semi-transparent CPK representations. Other residues involved in the interactions between TM2 and TM4’ in the adjacent subunit are shown in stick models. (**F**) Subunit interfaces at the CTDs of the three CALHM channels. The center CTDs are colored red, and the two adjacent CTDs are colored purple and green. The conserved residues among the CALHM channels are shown in stick models and semi-transparent CPK representations. Other residues involved in interactions between adjacent CTDs are shown in stick models. Hydrogen bonds and salt bridges are shown as black dashed lines.

Similar to OlCALHM1, both the TMD and CTD are involved in the subunit interactions in HsCALHM2 and CeCLHM-1 (Fig. 4D–F). At the TMD, TM2 of one subunit and TM4’ of the neighboring subunit form extensive hydrophobic interactions (Fig. 4D, E). Particularly, two Pro residues on TM2 and TM3, and two Trp residues on TM3 and TM4’ form van der Waals interactions, resulting in the tight contacts of the three TM helices (TM2, TM3, and TM4’), which are conserved among the three CALHM structures (Figs. 4D, E and S7). Structural comparisons of the three CALHM homologs show the different relative positions and orientations of TM2 relative to TM4’ of the neighboring subunits (Fig. 4D). The subunit interfaces between TM2 and TM4’ in OlCALHM1 are formed by tighter hydrophobic interactions, relative to those in HsCALHM2 and CeCLHM-1: The distances between the Cα atoms of the intracellular ends in TM2 and TM4’ in OlCALHM1, HsCALHM2, and CeCLHM-1 are 14.4, 17.9, and 16.8 Å, respectively (Fig. 4D). These subtle differences in the subunit interfaces at the TMD are likely to be associated with the different oligomeric assemblies. At the CTD in the three CALHM structures, the neighboring CTHs entangle together and form stacking interactions between the conserved aromatic residues: Phe and His residues in one subunit, a Phe residue in the adjacent subunit, and two Tyr residues from the adjacent subunit on the opposite side (Figs. 4F and S7). Taken together, the structural comparisons among the three CALHM homologs reveal both structural conservation (TM topology) and divergence (structures of CTD and oligomeric assembly).

## Oligomeric assembly mechanism of CALHM channels

The three different oligomeric assembly structures of the CALHM homologs probably arise from the slight variations in the inter-subunit interactions formed at the TMD and CTD. To investigate whether the TMD or CTD defines the oligomeric stoichiometry, we constructed a chimeric channel, consisting of the TMD from OlCALHM1 and the CTD from HsCALHM2 (named ‘OlCALHM1c’). The purified OlCALHM1c was exchanged into amphipol PMAL-C8 and subjected to the cryo-EM analysis. Without imposing any symmetry, the 3D classification provided cryo-EM maps of two different oligomeric states, the 8-mer and the 9-mer, but no 11-mer structure was obtained (Figs. 5A, S8). We finally determined the cryo-EM structures of the 8-mer and the 9-mer, both at resolutions of 3.4 Å (Figs. 5A, B, S8 and Table S1). The dimensions of the 8-mer and 9-mer structures of OlCALHM1c are consistent with the 8-mer and 9-mer structures of OlCALHM1 and CeCLHM-1, respectively (Fig. 5B). Since the densities of the CTDs are obscure in both the 8-mer and 9-mer EM maps (Fig. 5A), we did not model the entire CTD of the 8-mer and the side chains at the CTD of the 9-mer (Fig. 5B). By contrast, the EM density maps corresponding to the TMD are clearly resolved in both structures, allowing the identification of the residues in the TMD (Fig. 5B). The TMD structures of the respective subunits are almost identical between the 8-mer and the 9-mer (RMSD of 0.55 Å over 184 Cα atoms) (Fig. 5C).

**Fig. 5:**
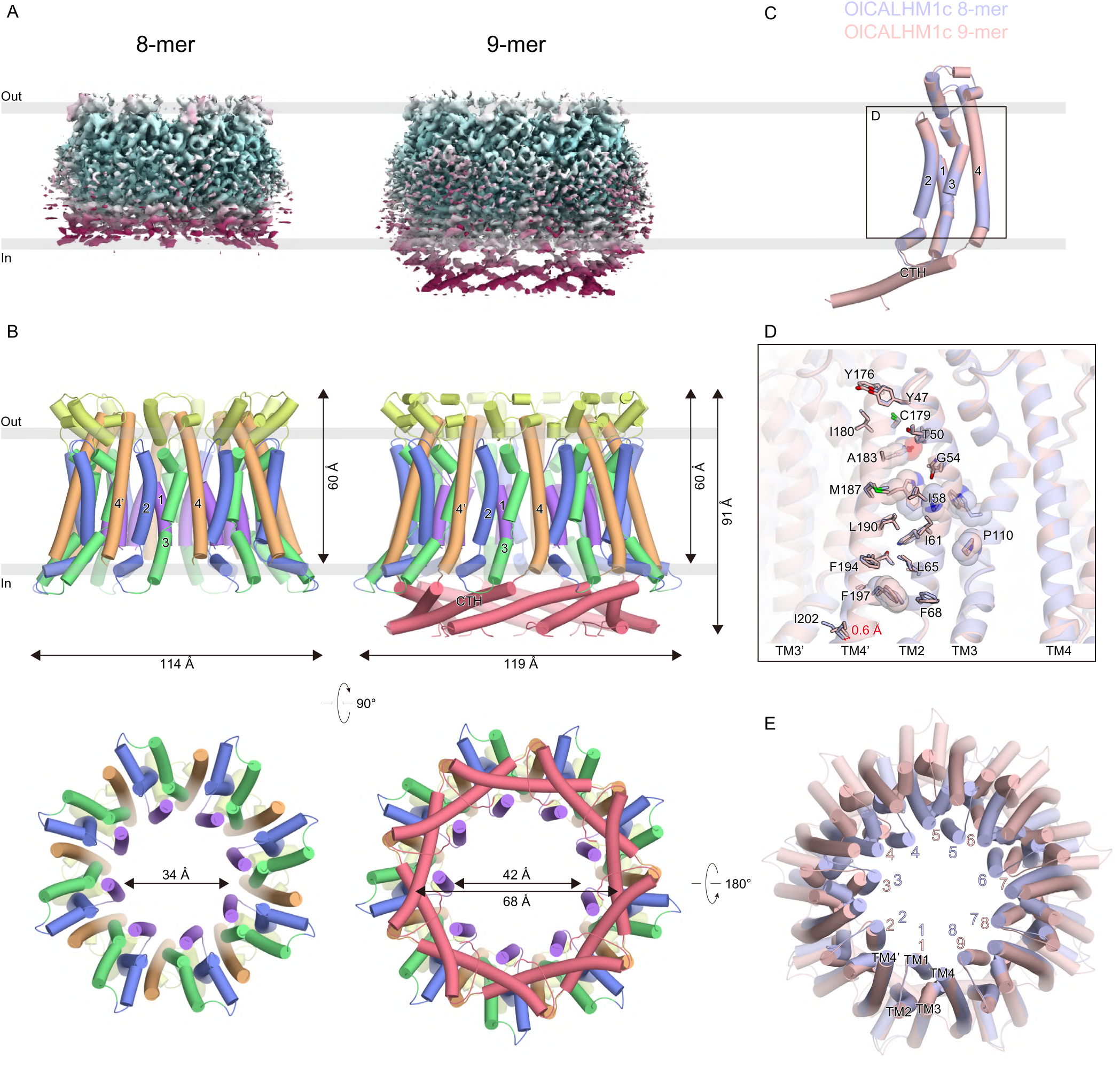
Structures and oligomerization states of the chimeric CALHM proteins. (**A**) EM density maps of the OlCALHM1c 8-mer (left) and 9-mer (right), colored according to the local resolution (blue: high resolution to red: low resolution), estimated by RELION. (**B**) Overall structures of the OlCALHM1c 8-mer (left) and 9-mer (right), viewed from the membrane side, colored as in Fig. 1D. (**C**) Superimposition of the OlCALHM1c 8-mer and 9-mer, viewed from the membrane side. The 8-mer and 9-mer structures are colored pale blue and pale pink, respectively. (**D**) Close-up view of the superimposition of the OlCALHM1c 8-mer and 9-mer, colored as in Fig. 5C. The conserved residues among the CALHM channels are shown in stick models and semi-transparent CPK representations. The red arrow indicates the different orientations of TM4’ in the adjacent subunit between the 8-mer and 9-mer structures. (**E**) Superimposition of the OlCALHM1c 8-mer and 9-mer, viewed from the extracellular side, colored as in Fig. 5C. The two structures are superimposed based on TM2 of subunit 1.

The subunit interfaces at the TMD of OlCALHM1c are almost the same as those of OlCALHM1_EM_, formed by hydrophobic interactions between TM2 of one subunit and TM4’ (Fig. 5D). Consistently, the 8-mer structure can be well superimposed with OlCALHM1_EM_, with the same TMD element. On the other hand, superimposition of the 8-mer and the 9-mer based on a subunit shows a 0.6 Å displacement of the Cα atoms of Ile202 on the intracellular end of TM4’ (Fig. 5D). The substitution of the CTD probably allowed this subtle difference of TM4’ at the TMD subunit interfaces, resulting in the 9-mer assembly (Fig. 5B, E). These observations indicate that the assembly stoichiometry is mainly defined by the subunit interactions at the TMD (*e.g.,* the TMD of OlCALHM1 forms the 8-mer or the 9-mer). This notion suggests that the diverse oligomeric states in CALHM channels are generally determined by the subunit interactions at the TMD. Although the three CALHM homologs adopt similar TM topologies, the TMD of OlCALHM1 has low sequence identities with those of HsCALHM2 (29%) and CeCLHM-1 (23%). These sequence differences within the TMDs among the CALHM family members are likely to be reflected in their diverse oligomeric assemblies and eventually in the different diameters of the channel pore, which may be associated with the different specificities of the released substrates at distinct expression sites (*5*), (*11*), (*14*).

## Discussion

In this study, we present the cryo-EM structures of three CALHM homologs and the two oligomeric states of the chimeric construct, revealing the ATP conduction and oligomeric assembly mechanisms. Notably, the present 2.66 Å-resolution OlCALHM1 structure allowed the accurate modeling of the NTH and TM1 (Fig. S4A), which form the pore architecture in the Ca^2+^-free open state (Fig. 3A, B). The analysis of the ΔNTH mutants indicates that the NTH is involved in the channel conduction of the ATP (Fig. 3D). Although the NTH and TM1 are disordered in our HsCALHM2 and CeCLHM-1 structures, the comparable arrangement of the TM helices among OlCALHM1, HsCALHM2, and CeCLHM-1 suggests that the NTH would similarly constitute the channel pores in the CALHM family members. The pore constriction by the NTH is also observed in gap junction channels (*16*), (*17*), indicating that this is a conserved structural feature of these oligomeric channels, despite the lack of sequence similarity among them.

The ΔNTH mutant showed almost no ATP-release activity upon Ca^2+^-binding, similar to the wild-type (Fig. 3E), suggesting that other structural components are associated with the channel gating in OlCALHM1 (Fig. 6A, B). In the OlALHM1_EM_ structure, TM1 protrudes towards the channel pore, as compared to the other TMs (Fig. 1D). Recently, the cryo-EM structures of HsCALHM2 were reported in the open and inhibited states at 3.3 and 2.7 Å resolutions, respectively (*22*). These structures showed channel pore closing by the swing motion of TM1 by about 60° towards the pore axis, driven by the binding of the inhibitor RUR at the base of the TM segment. Therefore, these structures suggest the large rearrangement of TM1 upon the channel gating (*22*). Given the structural and sequence similarities between OlCALHM1 and HsCALHM2 (Figs. 4B, C and S7), TM1 may undergo a large structural movement and block the pore upon Ca^2+^ binding in OlCALHM1 (Fig. 6A, B).

**Fig. 6:**
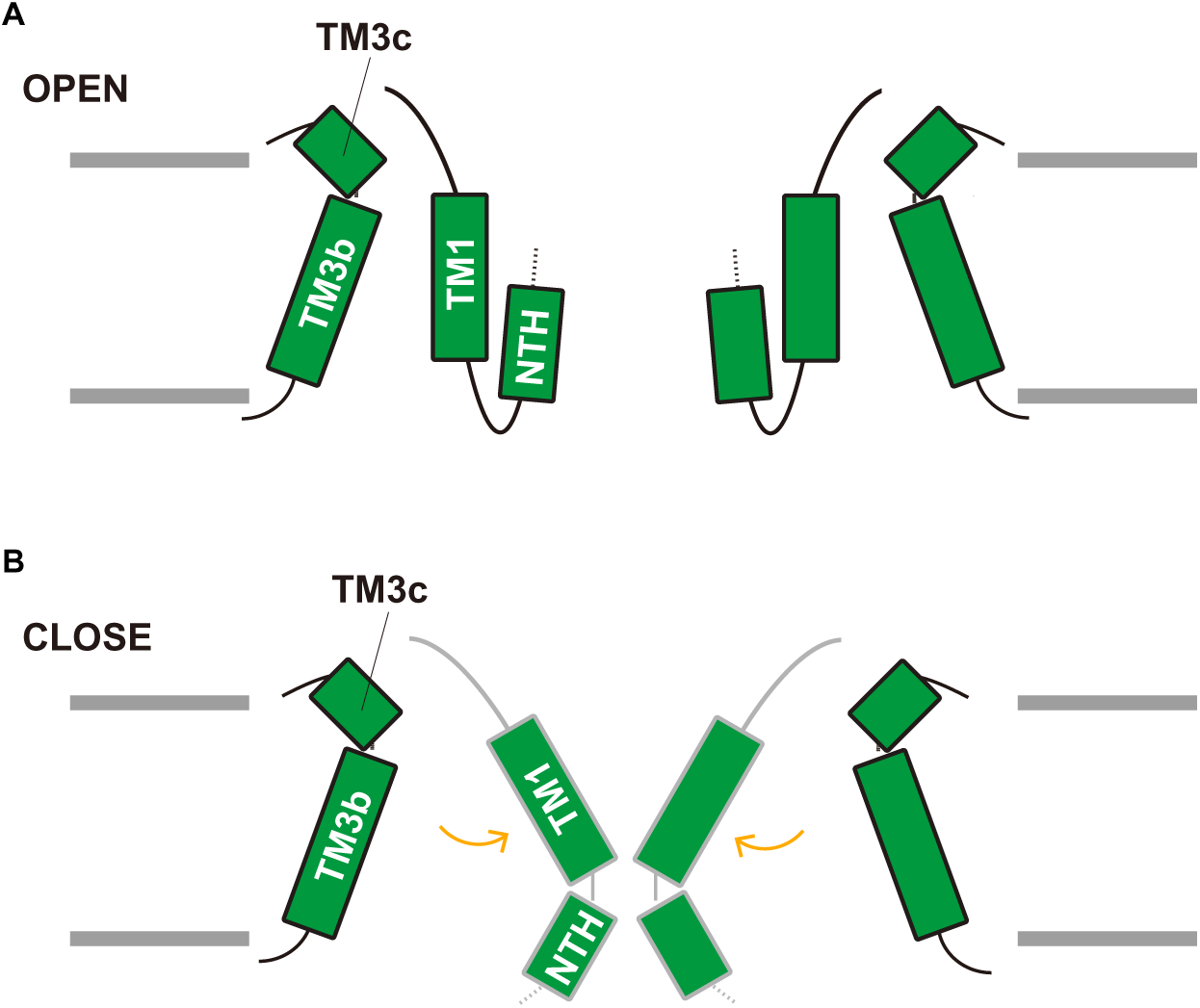
Insight into channel gating. (**A** and **B**) Schematic model of the Ca^2+^-dependent gating mechanism. In the Ca^2+^-free open state, TM1 forms the stable interaction with TM3 and moves perpendicularly to the membrane. The NTH forms the constriction site in the pore (A). In the Ca^2+^-bound closed state, TM1 blocks the pore (B).

For the subunit assembly of CALHM family proteins, the interactions at the TMD are mostly important for the oligomeric stoichiometry. The hydrophobic residues involved in the subunit interactions between TM2–TM4’ at the TMD are conserved in the CALHM1 orthologs, except for Met187 and Phe194 (Fig. S1A–C), implying a similar assembly number of HsCALHM1. In taste bud cells, CALHM1 and CALHM3 form a functionally heteromeric channel with fast activation (*12*). HsCALHM1 and HsCALHM3 have sequence identities of 47% overall and 54% within the TMD (Fig. S9), which are relatively higher as compared to the sequence identity between HsCALHM1 and HsCALHM2 (30% over the full-length and 33% within the TMD). Moreover, the residues that are probably involved in the subunit assembly are conserved in both HsCALHM1 and HsCALHM3, implying that they can form a heterooctamer, although the different residues of the CTD could slightly modify this assembly. Overall, our current study provides the ATP conduction and diverse assembly mechanisms of the CALHM family proteins.

## Acknowledgements

We thank T. Nakane for assistance with the single-particle analysis. We also thank the staff scientists at the University of Tokyo’s cryo-EM facility, especially K. Kobayashi, H. Yanagisawa, A. Tsutsumi, M. Kikkawa, and R. Danev.

## Authors’ contributions

K.D. prepared the cryo-EM sample of OlCALHM1 and performed the liposome assay. K.D. and T.K. collected and processed the cryo-EM data, and built the structure of OlCALHM1. T.K. prepared the cryo-EM sample of the CALHM chimera, collected and processed the cryo-EM data, and built the structures of HsCALHM2, CeCLHM-1, and the CALHM chimera. W.S. initially screened the CALHM homologs, established the expression and purification procedure, and prepared the cryo-EM sample of HsCALHM2. M.H. screened and prepared the cryo-EM sample of CeCLHM-1, collected and processed the cryo-EM data, and built the structure of CeCLHM-1. T.N. assisted with the data collection and processing. K.Y. assisted with the data processing and structure refinement. K.N. and A.T. measured the ATP-release activities of the CALHM proteins. K.D., T.K., W.S., and O.N. mainly wrote the manuscript. A.T. and O.N. supervised the research.

## Competing interests

M.H. is a graduate student at Mitsubishi Tanabe Pharma Corporation and is supported by the company with non-research funds. The company has no financial or other interest in this research.

**Data and materials availability:**

## Supplementary Materials

### Materials and Methods

#### Expression and purification of OlCALHM1, HsCALHM2, and the chimeric construct OlCALHM1c

The DNAs encoding the *Oryzias latipes* CALHM1 channel (UniProt ID: H2MCM1) and the *Homo sapiens* CALHM2 (UniProt ID: Q9HA72) were codon optimized for eukaryotic expression systems, synthesized (Genewiz, Inc.), and served as the templates for subcloning. To improve the expression and thermostability, the C-terminal residues starting from Trp296 were truncated (OlCALHM1_EM_). The chimeric construct consists of the TMD in OlCALHM1 (Met1–Ala209) followed by the CTD in HsCALHM2 (Ser213–Ser323). These DNA fragments were ligated into a modified pEG BacMam vector, with a C-terminal green fluorescent protein (GFP)-His_8_ tag and a tobacco etch virus (TEV) cleavage site, and were expressed in HEK293S GnTI^−^ (N-acetylglucosaminyl-transferase I-negative) cells (American Type Culture Collection, catalog no. CRL-3022) (*24*). Cells were collected by centrifugation (8,000 × *g*, 10 min, 4°C). For the cells expressing OlCALHM1c, the cells were disrupted by probe sonication and the membrane fraction was collected by ultracentrifugation (186,000 × g, 1 h, 4°C). The cells or the membrane fraction were solubilized for 1 h in buffer containing 50 mM Tris, pH 8.0, 150 mM NaCl, 2 mM dithiothreitol (DTT), 1% DDM (Calbiochem), and 0.2% cholesterol hemisuccinate (CHS). The mixture was centrifuged at 40,000g for 20 min, and the supernatant was incubated with CNBr-Activated Sepharose 4 Fast Flow beads (GE Healthcare) coupled with an anti-GFP nanobody (GFP enhancer or GFP minimizer) (*25*) for 2 h at 4°C. The protein-bound resins were washed with gel filtration buffer (50 mM Tris, pH 8.0, 150 mM NaCl, 2 mM dithiothreitol) containing 0.05% GDN for OlCALHM1_EM_ and OlCALHM1c or 0.06% digitonin for HsCALHM2, and further incubated overnight with TEV protease. The flow through was concentrated using a centrifugal filter unit (Merck Millipore, 100 kDa molecular weight cutoff), and separated on a Superose 6 Increase 10/300 GL column (GE Healthcare) equilibrated with gel filtration buffer. The peak fractions of the protein were pooled.

#### Expression and purification of CeCLHM-1

The *Caenorhabditis elegans* calcium homeostasis modulator protein (CeCLHM-1, Uniprot: Q18593) cDNA was synthesized, codon-optimized for expression in human cell lines, and cloned into the pcDNA3.4 vector. The CeCLHM-1 sequence was fused with a C-terminal His8 tag and enhanced green fluorescent protein (EGFP), followed by a tobacco etch mosaic virus (TEV) protease site. HEK293S GnTI^−^ cells were transiently transfected at a density of 2.0 × 10^6^ cells ml^−1^, with the plasmids and FectoPRO (Polyplus). Approximately 360 μg of the CeCLHM-1 plasmid were premixed with 360 μL FectoPRO reagent in 80 mL of fresh FreeStyle 293 medium, for 10–20 min before transfection. For transfection, 80 ml of the mixture was added to 0.8 L of cell culture and incubated at 37°C in the presence of 8% CO_2_. After 18–20 h, 2.2 mM valproic acid was added, and the cells were further incubated at 30°C in the presence of 8% CO_2_ for 48 h. The cells were collected by centrifugation (3,000 × g, 10 min, 4°C) and disrupted by Dounce homogenization in hypotonic buffer (50 mM HEPES-NaOH (pH 7.5), 10 mM KCl, 0.04 mg ml^−1^ DNase I, and protease inhibitor cocktail). The membrane fraction was collected by ultracentrifugation (138,000 × g, 1 h, 4°C), and solubilized for 1 h at 4°C in buffer (50 mM HEPES-NaOH (pH 7.5), 300 mM NaCl, 1.5% (w/v) N-dodecyl β-D-maltoside (DDM), and 0.15% (w/v) cholesterol hemisuccinate (CHS)). After ultracentrifugation (138,000 × g, 30 min, 4°C), the supernatant was incubated with AffiGel 10 (Bio-Rad) coupled with a GFP-binding nanobody, for 2 h at 4°C. The resin was washed five times with 3 CVs of wash buffer (50 mM HEPES-NaOH (pH 7.5), 300 mM NaCl, and 0.06% glyco-diogenin (GDN)), and gently suspended overnight with TEV protease to cleave the His8-EGFP tag. After the TEV protease digestion, the flow-through was pooled, concentrated, and purified by size-exclusion chromatography on a Superose 6 Increase 10/300 GL column (GE Healthcare), equilibrated with SEC buffer (20 mM HEPES-NaOH (pH 7.5), 150 mM NaCl, and 0.06% GDN). The peak fractions of the protein were pooled.

#### Grid preparation

The purified OlCALHM1_EM_ protein in GDN was concentrated to 5 mg ml^−1^. For cryo-EM analyses of HsCALHM2, CeCLHM-1, and OlCALHM1c, the detergents were exchanged with amphipols. The purified HsCALHM2 in digitonin, CeCLHM-1 in GDN, and OlCALHM1c in GDN were mixed with amphipol PMAL-C8 (Anatrace) at a 1:100 ratio (w/w) for 2–2.5 h at 4°C. Detergents were removed using Bio-Beads SM-2 (Bio-Rad, 100 mg per 1 ml mixture) for 2–2.5 h at 4°C. After removal of the Bio-Beads, the samples were concentrated and subjected to size-exclusion chromatography on a Superose 6 Increase 10/300 GL column (GE Healthcare), equilibrated with buffer (20 mM Tris-HCl, pH 8.0, 150 mM NaCl, 2 mM dithiothreitol). The peak fractions containing HsCALHM2, CeCLHM-1, and the OlCALHM1c in PMAL-C8 were concentrated to 1.6, 2.2, and 1 mg ml^−1^, respectively. Portions (3 µl) of the concentrated samples were applied to glow-discharged R1.2/1.3 Cu/Rh 300 mesh grids (Quantifoil). The grids were subsequently blotted (4 s for OlCALHM1_EM_, 8 s for HsCALHM2, 4 s for CeCLHM-1, and 4 s for OlCALHM1c) and vitrified using a Vitrobot Mark IV (FEI) under 4°C and 100% humidity conditions.

#### EM image acquisition and data processing

For OlCALHM1_EM_ and OlCALHM1c, data collections were performed on an FEI Titan Krios (FEI) electron microscope, operating at an acceleration voltage of 300 kV and equipped with a BioQuantum K3 imaging filter and a K3 direct electron detector (Gatan). EM images were acquired at a nominal magnification of 105,000 ×, corresponding to a physical pixel size of 0.83 Å, using the SerialEM software (*26*). For OlCALHM1_EM_, movies were dose fractionated to 54 frames at a dose rate of 14.9 e^−^ pixel^−1^ per second, resulting in a total accumulated dose of 59.4 e^−^ Å^−2^. For OlCALHM1c, movies were dose fractionated to 48 frames at a dose rate of 14.0 e^−^ pixel^−1^ per second, resulting in a total accumulated dose of 50 e^−^ Å^−2^. For HsCALHM2 and CeCLHM-1, data collections were performed on a 300 kV Titan Krios electron microscope equipped with a Falcon III direct electron detector. EM images were acquired at a nominal magnification of 96,000 ×, corresponding to a calibrated pixel size of 0.8346 Å pixel^−1^, using the EPU software. For HsCALHM2, movies were dose fractionated to 60 frames at a dose rate of 0.95 e^−^ pixel^−1^ per second, resulting in a total accumulated dose of 60 e^−^ Å^−2^. For CeCLHM-1, movies were dose fractionated to 48 frames at a dose rate of 0.7 e^−^ pixel^−1^ per second, resulting in a total accumulated dose of 50 e^−^ Å^−2^. For all datasets, the movie frames were aligned in 4 × 4 patches and dose-weighted using RELION (*27*). CTF estimation was performed by CTFFIND 4.1 (*28*).

For the OlCALHM1_EM_ dataset, a total of 1,231,992 particles were extracted from 3,075 micrographs in 3.55 Å pixel^−1^ using AutoPick in RELION-3.0 (*27*), applying the two-dimensional class averages of reference-free auto-picked particles based on a Laplacian-of-Gaussian (LoG) filter as templates. The initial model was generated in RELION. The particles were passed to RELION to calculate the two-dimensional classification, refinement, and post-processing. Since some classes of the two-dimensional classification exhibited the density for eight subunits, 137,146 particles in 3 good classes were re-extracted in the original pixel size of 0.83 Å pixel^−1^ and refined in C8 symmetry. The particles were further cleaned using 3D classification, and 102,109 good particles were subsequently subjected to micelle subtraction, polishing (*29*), and CTF refinement. The overall gold-standard resolution calculated at the Fourier shell correlation (FSC = 0.143) (*30*) was 2.66 Å.

For the HsCALHM2 dataset, a total of 946,353 particles were initially extracted from 2,110 micrographs, with a binned pixel size of 3.52 Å pixel^−1^, as described above. These particles were subjected to several rounds of 2D and 3D classifications, resulting in the best class from the 3D classification, which contained 365,116 particles. These particles were then re-extracted with 1.40 Å pixel^−1^, and subsequently subjected to 3D refinement with C11 symmetry, polishing, CTF refinement, and micelle subtraction. The final 3D refinement with C11 symmetry and postprocessing yielded a density map with an overall resolution of 3.51 Å at a gold-standard FSC = 0.143. The image processing workflow is described in Fig. S5.

For the CeCLHM-1 dataset, a total of 832,750 particles were initially extracted from 2,906 micrographs, with a binned pixel size of 3.01 Å pixel^−1^, as described above. These particles were subjected to several rounds of 2D classification, resulting in the best classes containing 261,169 particles. These particles were re-extracted with 1.5 Å pixel^−1^, and subjected to 3D classification, resulting in the best classes containing 65,887 particles. The particles were subsequently subjected to 3D refinement with C9 symmetry, polishing, and CTF refinement. The final 3D refinement with C9 symmetry and postprocessing yielded a density map with an overall resolution of 3.6 Å at a gold-standard FSC = 0.143. The image processing workflow is described in Fig. S6.

For the OlCALHM1c dataset, a total of 1,689,949 particles were initially extracted from 2,808 micrographs, with a binned pixel size of 3.39 Å pixel^−1^, as described above. These particles were subjected to several rounds of 2D and 3D classifications, resulting in the two distinct classes from the 3D classification, which represented the 8-mer and 9-mer structures. The 8-mer and 9-mer classes contained 233,566 and 227,781 particles, respectively. These particles were separately re-extracted with 1.41 Å pixel^−1^, and subsequently subjected to 3D refinement with C8 or C9 symmetry, respectively. The final 3D refinement and postprocessing of the two classes yielded density maps with overall resolutions of 3.4 Å in both cases, at a gold-standard FSC = 0.143. The image processing workflow is described in Fig. S8.

#### Model building and refinement

The model of OlCALHM1_EM_ was built *de novo* into the density map using COOT (*31*) and rebuilt using Rosetta (*32*). The model was again rebuilt using COOT with multiple rounds of real-space refinement in PHENIX (*33*), (*34*), with secondary structure restraints. The models of HsCALHM2 and CeCLHM-1 were built using COOT with multiple rounds of real-space refinement in PHENIX, with secondary structure restraints. For the OlCALHM1c 8-mer and 9-mer, the TMD of OlCALHM1_EM_ were modeled into the density map by rigid-body fitting and modified using COOT with multiple rounds of real-space refinement in PHENIX, with secondary structure restraints.

For cross-validation, the final models were randomly shaken and refined against one of the half maps generated in RELION. Fourier shell correlation (FSC) curves were calculated between the refined model and the half map 1, and between the refined model and the half map 2 (Figs. S3, S5, S6, and S8) (*35*). The statistics of the model refinement and validation are summarized in Table S1. Figures of the density maps and molecular graphics were prepared using CueMol (http://www.cuemol.org) and UCSF Chimera (*36*).

#### Bioluminescent ATP release assay

The extracellular ATP concentration was measured by the luciferin-luciferase reaction as previously reported (*21*). HeLa cells seeded at a density of 1.2 × 10^4^ cells well^−1^ on 96-well cell culture plates (Corning Costar, Corning, NY, USA) were transfected with 0.6 µg pIRES2.AcGFP1 encoding WT or mutant OlCALHM1 cDNA or the empty vector using Lipofectamine 3000 (Thermo Fisher Scientific), according to the manufacturer’s instruction, and incubated for 4 h at 37 °C. Subsequently, transfection media were replaced with fresh culture media, and cells were incubated at 27 °C for 4 h before being tested. In experiments shown in Fig. S2A–C, cells were transfected with 0.2 µg pIRES2.AcGFP1 encoding HsCALHM1, HsCALHM2, OlCALHM1, or CeCLHM-1 cDNA or the empty vector, and incubated in the transfection media for 4 h at 37 °C and then in fresh culture media for 20 h at 27 °C. Cells were then washed twice with the standard bath solution and incubated in 100 µL of the standard bath solution for another 1 h at room temperature. Immediately after 75 µl of the bath solution was replaced with an equal volume of the Ca^2+^-free bath solution to make final [Ca^2+^]_o_ ∼17 nM, 10 µl of the luciferin-luciferase solution (FL-AAM and FL-AAB, Sigma, St. Louis, MO, USA) was added to each well and the plate was placed in a microplate luminometer (Centro LB960, Berthold Technologies, Bad Wildbad, Germany). Luminescence was measured every 2 min at 25 °C. The extracellular ATP concentration was calculated from a standard curve created in each plate. The standard bath solution contained (in mM): 150 NaCl, 5 KCl, 2 CaCl_2_, 1 MgCl_2_, 10 HEPES, and 10 glucose, pH 7.4 adjusted with NaOH. The Ca^2+^-free bath solution contained (in mM): 150 NaCl, 5 KCl, 5 EGTA, 1 MgCl_2_, 10 Hepes, and 10 glucose, pH 7.4 adjusted with NaOH.

#### Preparation of proteoliposomes

Lipids from chloroform stocks, mixed at a 2:1 weight ratio of 1-palmitoyl-2-oleoyl-sn-glycero-3-phosphoglycerol (Avanti) with 12% egg phosphatidylcholine:1-palmitoyl-2-oleoyl-sn-glycero-3-phosphoglycerol (Avanti), were dried and solubilized in a solution containing 100 mM KCl, 0.1 mM EGTA, 2.3% n-octyl-β-D-glucopyranoside, and 25 mM HEPES, pH 7.6. Solubilized OlCALHM1_EM_ was added to the lipid-detergent mixture at a 50:1 lipid to protein ratio. After a short incubation, Bio-Beads SM-2 (Bio-Rad) were added and mixed at 4°C overnight. To confirm the reconstitution, after ultracentrifugation (125,000g, 15 min, 4°C) and solubilization in buffer (50 mM Tris–HCl, pH 8.0, 150 mM NaCl, 1.0% DDM, and 0.2% CHS) at 4°C for 1 h, the proteoliposomal solutions were subjected to SDS-PAGE.

#### ATP transport assay

Liposomes and proteoliposomes were loaded with 1 mM ATP and sonicated for 30 s, using a Bioruptor (CosmoBio). After removal of the extraliposomal ATP by chromatography on Sephadex G-50 fine resin (GE Healthcare), 50 μl portions of the samples were combined with 50 μl of a mixture containing 45 nM luciferase (QuantiLum *Photinus pyralis* recombinant luciferase; Promega, Madison, WI), 1.2 nM luciferin (Nacalai Tesque), 1.0 mM EDTA, and 10 mM MgSO_4_. After 1h incubation with or without 0.4% Triton X-100, the luminescence was measured on an ARVO-X3 microplate reader (PerkinElmer). The amount of ATP retained in the liposomes was determined as the difference in the luminescence between total ATP and extraliposomal ATP that was accessible to luciferase without Triton X-100.

**Fig. S1: Amino acid sequence alignment of the CALHM1 orthologs.**

(**A**) Alignment of the amino-acid sequences of killifish CALHM1 (OlCALHM1, UniProt ID: H2MCM1), human CALHM1 (UniProt ID: Q8IU99), rat CALHM1 (UniProt ID: D4AE44), cat CALHM1 (UniProt ID: M3VYF2), chicken CALHM1 (UniProt ID: A0A1D5NWS1), and *Danio rerio* CALHM1 (UniProt ID: E7F2J4). Secondary structure elements for α-helices are indicated by cylinders. Conservation of the residues is indicated as follows: red panels for completely conserved; red letters for partly conserved; and black letters for not conserved. (**B** and **C**) Conservation of the residues of CALHM1 family members. The sequence conservation among 384 CALHM1 orthologs was calculated using the ConSurf server (http://consurf.tau.ac.il), and is colored from cyan (low) to maroon (high). The residues constituting the subunit interface are indicated by stick models. They are conserved in the CALHM1 orthologs, except for Met187 and Phe194, which are exposed to the membrane environment.

**Fig. S2: Functional analyses of the CALHM proteins.**

(**A** and **B**) Time courses of extracellular ATP levels due to release from HeLa cells transfected with the empty vector (mock), OlCALHM1, HsCALHM1, HsCALHM2, or CeCLHM-1 following exposure to essentially zero [Ca^2+^]_o_ (17 nM), in the absence (A) or presence (B) of RUR (20 μM). Data are displayed as means ± S.E.M. (standard error of the mean) (n = 8). (**C**) Time courses of extracellular ATP levels due to release from the empty vector (mock), full-length OlCALHM1, or OLCALHM1_EM_ transfected HeLa cells, following exposure to essentially zero [Ca^2+^]_o_ (17 nM). Data are displayed as means ± S.E.M. (n = 8). (**D**) Permeability of the purified OlCALHM1_EM_ protein to ATP. Liposomes and OlCALHM1_EM_ proteoliposomes loaded with 1 mM ATP were fractionated on the size exclusion column to separate the free extraliposomal ATP from the ATP retained inside. The retained ATP was measured by luminescence using a luciferin/luciferase assay, as the difference between total ATP (inside plus outside, measured after the addition of Triton X-100 to 0.4%) and extraliposomal ATP (before Triton X-100). Data are displayed as the means ± S.E.M. (n = 3) of the ATP retained/total counts.

**Fig. S3: Cryo-EM analysis of OlCALHM1.**

(**A**) Representative cryo-EM image of OlCALHM1, recorded on a 300 kV Titan Krios electron microscope equipped with a K3 camera. (**B**) Image processing workflow of the single particle analysis. (**C**) Angular distribution. (**D**) Fourier shell correlation curve of two half-maps. (**E**) Map-to-model correlation curves.

**Fig. S4: Atomic model of OlCALHM1 in the density maps.**

(**A**) Cryo-EM density and atomic model of each segment of OlCALHM1_EM_. All maps are contoured at 4.0 σ. (**B**) Structural comparison of OlCALHM1_EM_ and Gap-like channels (connexin 26, orange; innexin, light green; and LRRC8, purple).

**Fig. S5: Cryo-EM analysis of HsCALHM2.**

(**A**) Representative cryo-EM image of HsCALHM2, recorded on a 300 kV Titan Krios electron microscope equipped with a Falcon III camera. (**B**) Image processing workflow of the single particle analysis. (**C**) Angular distribution. (**D**) Fourier shell correlation curve of two half-maps. (**E**) Local resolution analysis. (**F**) Map-to-model correlation curves.

**Fig. S6: Cryo-EM analysis of CeCLHM-1.**

(**A**) Representative cryo-EM image of CeCLHM-1, recorded on a 300 kV Titan Krios electron microscope equipped with a Falcon III camera. (**B**) Image processing workflow of the single particle analysis. (**C**) Angular distribution. (**D**) Fourier shell correlation curve of two half-maps. (**E**) Local resolution analysis. (**F**) Map-to-model correlation curves.

**Fig. S7: Amino acid sequence alignment of OlCALHM1, HsCALHM2, and CeCLHM-1.**

Alignment of the amino-acid sequences of killifish CALHM1 (OlCALHM1, UniProt ID: H2MCM1), human CALHM1 (UniProt ID: Q8IU99), human CALHM2 (UniProt ID: Q9HA72), and CeCLHM-1 (UniProt ID: Q18593). Secondary structure elements for α-helices are indicated by cylinders. Conservation of the residues is indicated as follows: red panels for completely conserved; red letters for partly conserved; and black letters for not conserved.

**Fig. S8: Cryo-EM analysis of the CALHM chimera.**

(**A**) Representative cryo-EM image of the CALHM chimera (OlCALHM1c), recorded on a 300 kV Titan Krios electron microscope equipped with a K3 camera. (**B**) Image processing workflow of the single particle analysis. (**C**) Angular distributions of the OlCALHM1c 8-mer (left) and 9-mer (right). (**D**) Fourier shell correlation curves of two half-maps of the OlCALHM1c 8-mer (left) and 9-mer (right). (**E**) Local resolution analysis of the OlCALHM1c 8-mer (left) and 9-mer (right). (**F**) Map-to-model correlation curves of the OlCALHM1c 8-mer (left) and 9-mer (right).

**Fig. S9: Amino acid sequence alignment of OlCALHM1, HsCALHM1, and HsCALHM3.**

Alignment of the amino-acid sequences of killifish CALHM1 (OlCALHM1, UniProt ID: H2MCM1), human CALHM1 (UniProt ID: Q8IU99), and human CALHM3 (UniProt ID: Q86XJ0). Secondary structure elements for α-helices are indicated by cylinders. Conservation of the residues is indicated as follows: red panels for completely conserved; red letters for partly conserved; and black letters for not conserved.

**Table S1:** Cryo-EM data collection, processing, model refinement, and validation statistics.

|  | OICALHM1 | HsCALHM2 | CeCLHM-1 | Chimeric<br>construct (8-mer) | Chimeric<br>construct (9-mer) |
| --- | --- | --- | --- | --- | --- |
| EMDB ID | XXXX | XXXX | XXXX | XXXX | XXXX |
| PDB ID | XXXX | XXXX | XXXX | XXXX | XXXX |
| **Data collection and processing** |  |  |  |  |  |
| Microscope | Titan Krios G3i | Titan Krios G3i | Titan Krios G3i | Titan Krios G3i |  |
| Detector | Gatan K3 camera | FEI Falcon III | FEI Falcon III | Gatan K3 camera |  |
| Nominal magnification | 105,000 | 96,000 | 96,000 | 105,000 |  |
| Voltage (kV) | 300 | 300 | 300 | 300 |  |
| Electron exposure (e<sup>-</sup> Å<sup>-2</sup>) | 59 | 60 | 50 | 50 |  |
| Nominal defocus range (μm) | −0.8 to −1.6 | −0.75 to −1.75 | −0.75 to −1.75 | −0.8 to −1.52 |  |
| Pixel size (Å pixel<sup>-1</sup>) | 0.83 | 0.8346 | 0.8346 | 0.83 |  |
| Symmetry imposed | C8 | C11 | C9 | C8 | C9 |
| Number of movies | 3,075 | 2,110 | 2,906 | 2,808 |  |
| Initial particle numbers | 1,231,992 | 946,353 | 832,750 | 1,689,949 |  |
| Final particle numbers | 102,109 | 365,116 | 65,887 | 233,566 | 227,781 |
| Map resolution (Å) | 2.66 | 3.51 | 3.60 | 3.4 | 3.4 |
| FSC threshold | 0.143 | 0.143 | 0.143 | 0.143 | 0.143 |
| Map sharpening *B* factor (Å<sup>2</sup>) | −61.6548 | −203.38 | −120.37 | −181.13 | −184.76 |
| **Model building and refinement** |  |  |  |  |  |
| Model composition |  |  |  |  |  |
| Protein atoms | 17,412 | 23,716 | 17,717 | 11,624 | 17,334 |
| Other atoms | 139 | 0 | 0 | 0 | 0 |
| R.M.S. deviations from ideal |  |  |  |  |  |
| Bond lengths (Å) | 0.009 | 0.004 | 0.008 | 0.007 | 0.008 |
| Bond angles (°) | 1.013 | 0.898 | 0.831 | 0.879 | 0.924 |
| **Validation** |  |  |  |  |  |
| Clashscore | 4.85 | 3.00 | 2.57 | 2.21 | 5.28 |
| Rotamer outliers (%) | 2.23 | 0.00 | 0.94 | 0.63 | 0.49 |
| Ramachandran plot |  |  |  |  |  |
| Favored (%) | 97.3 | 91.55 | 94.24 | 98.35 | 97.48 |
| Allowed (%) | 2.7 | 8.45 | 5.76 | 1.65 | 2.52 |
| Outlier (%) | 0.0 | 0.00 | 0.00 | 0.00 | 0.00 |

## References and Notes

1. G. Burnstock, Historical review: ATP as a neurotransmitter. Trends in Pharmacological Sciences. 27, 166–176 (2006).

2. M. P. Abbracchio, G. Burnstock, A. Verkhratsky, H. Zimmermann, Purinergic signalling in the nervous system: an overview. Trends in Neurosciences. 32, 19– 29 (2009).

3. A. Taruno, ATP release channels. International Journal of Molecular Sciences. 19, 808 (2018).

4. U. Dreses-Werringloer, J. C. Lambert, V. Vingtdeux, H. Zhao, H. Vais, A. Siebert, A. Jain, J. Koppel, A. Rovelet-Lecrux, D. Hannequin, F. Pasquier, D. Galimberti, E. Scarpini, D. Mann, C. Lendon, D. Campion, P. Amouyel, P. Davies, J. K. Foskett, F. Campagne, P. Marambaud, A Polymorphism in CALHM1 Influences Ca^2+^ Homeostasis, Aβ Levels, and Alzheimer’s Disease Risk. Cell. 133, 1149–1161 (2008).

5. A. Taruno, V. Vingtdeux, M. Ohmoto, Z. Ma, G. Dvoryanchikov, A. Li, L. Adrien, H. Zhao, S. Leung, M. Abernethy, J. Koppel, P. Davies, M. M. Civan, N. Chaudhari, I. Matsumoto, G. Hellekant, M. G. Tordoff, P. Marambaud, J. K. Foskett, CALHM1 ion channel mediates purinergic neurotransmission of sweet, bitter and umami tastes. Nature. 495, 223–226 (2013).

6. Z. Ma, W. T. Saung, J. K. Foskett, Action potentials and ion conductances in wild-type and CALHM1-knockout type II taste cells. Journal of Neurophysiology. 117, 1865–1876 (2017).

7. A. Taruno, I. Matsumoto, Z. Ma, P. Marambaud, J. K. Foskett, How do taste cells lacking synapses mediate neurotransmission? CALHM1, a voltage-gated ATP channel. BioEssays. 35, 1111–1118 (2013).

8. Z. Ma, A. P. Siebert, K. H. Cheung, R. J. Lee, B. Johnson, A. S. Cohen, V. Vingtdeux, P. Marambaud, J. K. Foskett, Calcium homeostasis modulator 1 (CALHM1) is the pore-forming subunit of an ion channel that mediates extracellular Ca^2+^ regulation of neuronal excitability. Proceedings of the National Academy of Sciences of the United States of America. 109, E1963–E1971 (2012).

9. Z. Ma, J. E. Tanis, A. Taruno, J. K. Foskett, Calcium homeostasis modulator (CALHM) ion channels. Pflugers Archiv European Journal of Physiology. 468, 395–403 (2016).

10. A. P. Siebert, Z. Ma, J. D. Grevet, A. Demuro, I. Parker, J. K. Foskett, Structural and functional similarities of calcium homeostasis modulator 1 (CALHM1) ion channel with connexins, pannexins, and innexins. Journal of Biological Chemistry. 288, 6140–6153 (2013).

11. J. Ma, X. Qi, C. Yang, R. Pan, S. Wang, J. Wu, L. Huang, H. Chen, J. Cheng, R. Wu, Y. Liao, L. Mao, F. Wang, Z. Wu, J. An, Y. Wang, X. Zhang, C. Zhang, Z. Yuan, Calhm2 governs astrocytic ATP releasing in the development of depression-like behaviors. Molecular Psychiatry. 23, 883–891 (2018).

12. Z. Ma, A. Taruno, M. Ohmoto, M. Jyotaki, J. C. Lim, H. Miyazaki, N. Niisato, Y. Marunaka, R. J. Lee, H. Hoff, R. Payne, A. Demuro, I. Parker, C. H. Mitchell, J. Henao-Mejia, J. E. Tanis, I. Matsumoto, M. G. Tordoff, J. K. Foskett, CALHM3 Is Essential for Rapid Ion Channel-Mediated Purinergic Neurotransmission of GPCR-Mediated Tastes. Neuron. 98, 547–561.e10 (2018).

13. M. Kashio, G. Wei-qi, Y. Ohsaki, M. A. Kido, A. Taruno, CALHM1/CALHM3 channel is intrinsically sorted to the basolateral membrane of epithelial cells including taste cells. Scientific Reports. 9, 2681 (2019).

14. J. E. Tanis, Z. Ma, P. Krajacic, L. He, K. J. Foskett, T. Lamitina, CLHM-1 is a functionally conserved and conditionally toxic Ca^2+^-permeable ion channel in *Caenorhabditis elegans*. Journal of Neuroscience. 33, 12275–12286 (2013).

15. J. E. Tanis, Z. Ma, J. K. Foskett, The NH_2_ terminus regulates voltage-dependent gating of CALHM ion channels. American Journal of Physiology Cell Physiology. 313, C173–C186 (2017).

16. S. Maeda, S. Nakagawa, M. Suga, E. Yamashita, A. Oshima, Y. Fujiyoshi, T. Tsukihara, Structure of the connexin 26 gap junction channel at 3.5 Å resolution. Nature. 458, 597–602 (2009).

17. A. Oshima, K. Tani, Y. Fujiyoshi, Atomic structure of the innexin-6 gap junction channel determined by cryo-EM. Nature Communications. 7, 13681 (2016).

18. G. Kasuya, T. Nakane, T. Yokoyama, Y. Jia, M. Inoue, K. Watanabe, R. Nakamura, T. Nishizawa, T. Kusakizako, A. Tsutsumi, H. Yanagisawa, N. Dohmae, M. Hattori, H. Ichijo, Z. Yan, M. Kikkawa, M. Shirouzu, R. Ishitani, O. Nureki, Cryo-EM structures of the human volume-regulated anion channel LRRC8. Nature Structural and Molecular Biology. 25, 797–804 (2018).

19. D. Deneka, M. Sawicka, A. K. M. Lam, C. Paulino, R. Dutzler, Structure of a volume-regulated anion channel of the LRRC8 family. Nature. 558, 254–259 (2018).

20. T. Kawate, E. Gouaux, Fluorescence-Detection Size-Exclusion Chromatography for Precrystallization Screening of Integral Membrane Proteins. Structure. 14, 673–681 (2006).

21. A. Taruno, H. Sun, K. Nakajo, T. Murakami, Y. Ohsaki, M. A. Kido, F. Ono, Y. Marunaka, Post-translational palmitoylation controls the voltage gating and lipid raft association of the CALHM1 channel. Journal of Physiology. 595, 6121–6145 (2017).

22. W. Choi, N. Clemente, W. Sun, J. Du, W. Lü, The structures and gating mechanism of human calcium homeostasis modulator 2. Nature. 576, 163–167 (2019).

23. O. S. Smart, J. G. Neduvelil, X. Wang, B. A. Wallace, M. S. P. Sansom, HOLE: A program for the analysis of the pore dimensions of ion channel structural models. Journal of Molecular Graphics. 14, 354–360 (1996).

24. A. Goehring, C. H. Lee, K. H. Wang, J. C. Michel, D. P. Claxton, I. Baconguis, T. Althoff, S. Fischer, K. C. Garcia, E. Gouaux, Screening and large-scale expression of membrane proteins in mammalian cells for structural studies. Nature Protocols. 9, 2574–2585 (2014).

25. A. Kirchhofer, J. Helma, K. Schmidthals, C. Frauer, S. Cui, A. Karcher, M. Pellis, S. Muyldermans, C. S. Casas-Delucchi, M. C. Cardoso, H. Leonhardt, K. P. Hopfner, U. Rothbauer, Modulation of protein properties in living cells using nanobodies. Nature Structural and Molecular Biology. 17, 133–139 (2010).

26. D. N. Mastronarde, Automated electron microscope tomography using robust prediction of specimen movements. Journal of Structural Biology. 152, 36–51 (2005).

27. J. Zivanov, T. Nakane, B. O. Forsberg, D. Kimanius, W. J. H. Hagen, E. Lindahl, S. H. W. Scheres, New tools for automated high-resolution cryo-EM structure determination in RELION-3. eLife. 7, e42166 (2018).

28. A. Rohou, N. Grigorieff, CTFFIND4: Fast and accurate defocus estimation from electron micrographs. Journal of Structural Biology. 192, 216–221 (2015).

29. J. Zivanov, T. Nakane, S. H. W. Scheres, A Bayesian approach to beam-induced motion correction in cryo-EM single-particle analysis. IUCrJ. 6, 5–17 (2019).

30. P. B. Rosenthal, R. Henderson, Optimal determination of particle orientation, absolute hand, and contrast loss in single-particle electron cryomicroscopy. Journal of Molecular Biology. 333, 721–745 (2003).

31. P. Emsley, B. Lohkamp, W. G. Scott, K. Cowtan, Features and development of Coot. Acta Crystallographica Section D: Biological Crystallography. 66, 486– 501 (2010).

32. R. Y. R. Wang, Y. Song, B. A. Barad, Y. Cheng, J. S. Fraser, F. DiMaio, Automated structure refinement of macromolecular assemblies from cryo-EM maps using Rosetta. eLife. 5, e17219 (2016).

33. P. V. Afonine, B. K. Poon, R. J. Read, O. V. Sobolev, T. C. Terwilliger, A. Urzhumtsev, P. D. Adams, Real-space refinement in PHENIX for cryo-EM and crystallography. Acta Crystallographica Section D: Structural Biology. 74, 531– 544 (2018).

34. P. D. Adams, P. V. Afonine, G. Bunkóczi, V. B. Chen, I. W. Davis, N. Echols, J. J. Headd, L. W. Hung, G. J. Kapral, R. W. Grosse-Kunstleve, A. J. McCoy, N. W. Moriarty, R. Oeffner, R. J. Read, D. C. Richardson, J. S. Richardson, T. C. Terwilliger, P. H. Zwart, PHENIX: A comprehensive Python-based system for macromolecular structure solution. Acta Crystallographica Section D: Biological Crystallography. 66, 213–221 (2010).

35. A. Brown, F. Long, R. A. Nicholls, J. Toots, P. Emsley, G. Murshudov, Tools for macromolecular model building and refinement into electron cryo-microscopy reconstructions. Acta Crystallographica Section D: Biological Crystallography. 71, 136–153 (2015).

36. E. F. Pettersen, T. D. Goddard, C. C. Huang, G. S. Couch, D. M. Greenblatt, E. C. Meng, T. E. Ferrin, UCSF Chimera - A visualization system for exploratory research and analysis. Journal of Computational Chemistry. 25, 1605–1612 (2004).

