## Supplementary figures and images for "Cryo-EM structures of calcium homeostasis modulator channels in diverse oligomeric assemblies"

### SupplementaryFigures

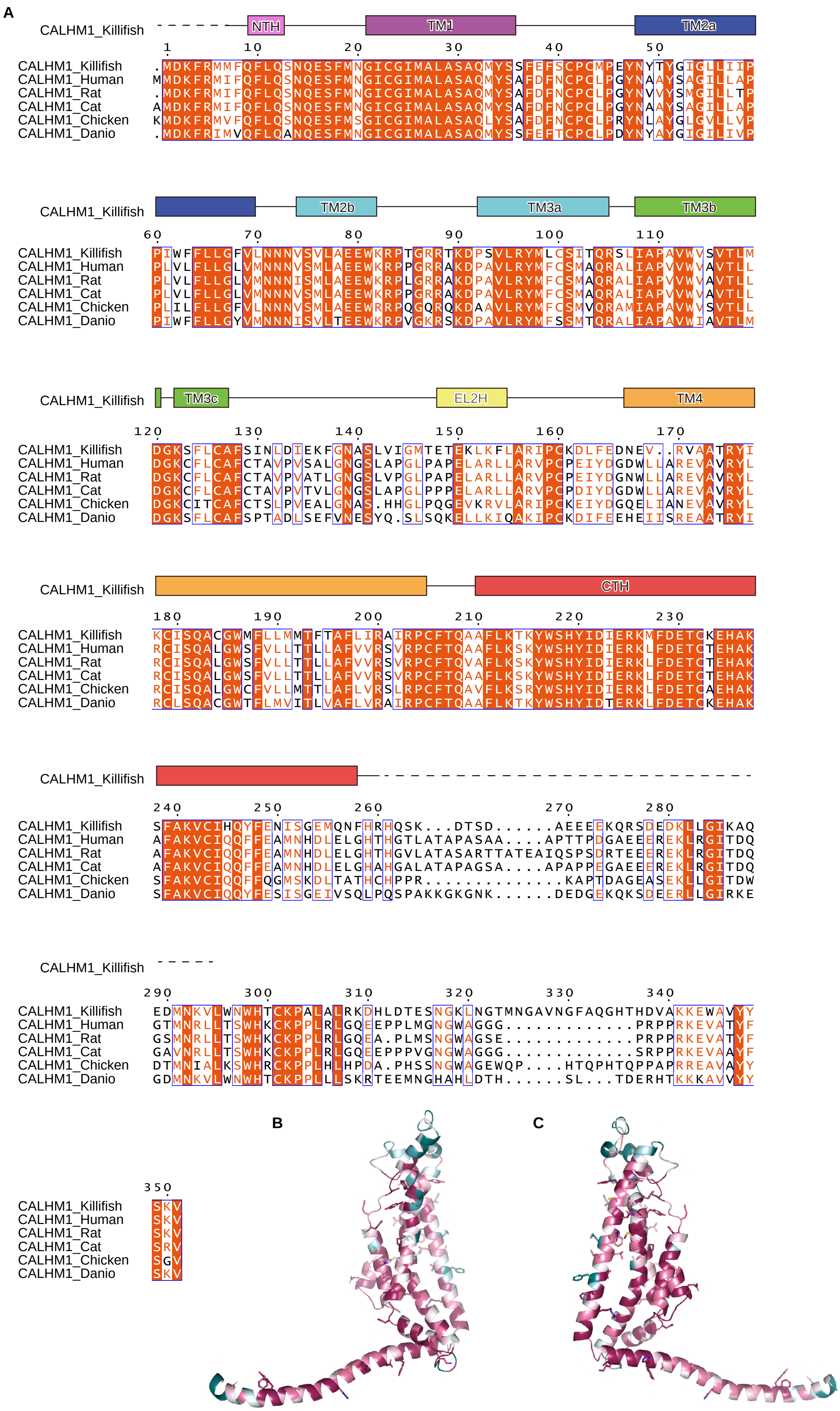

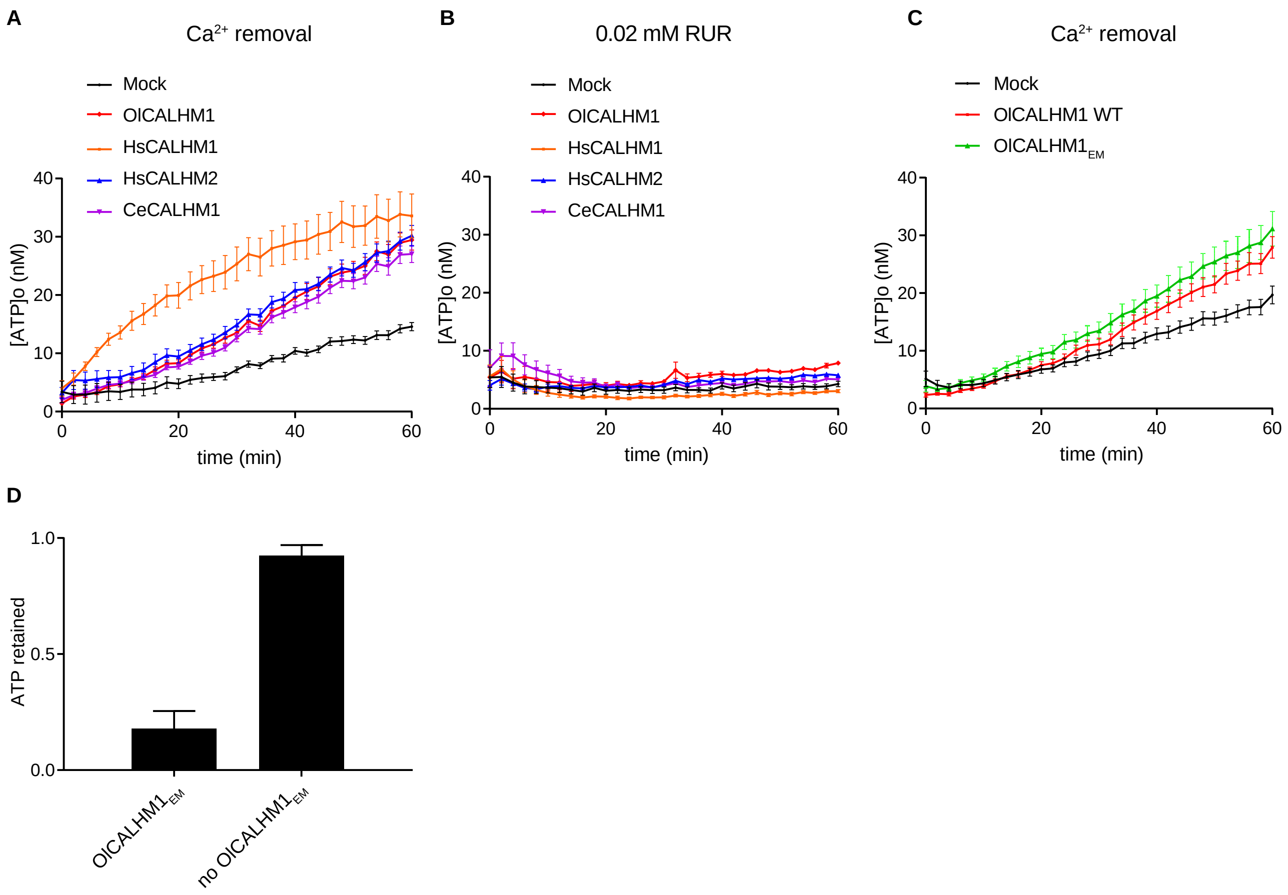

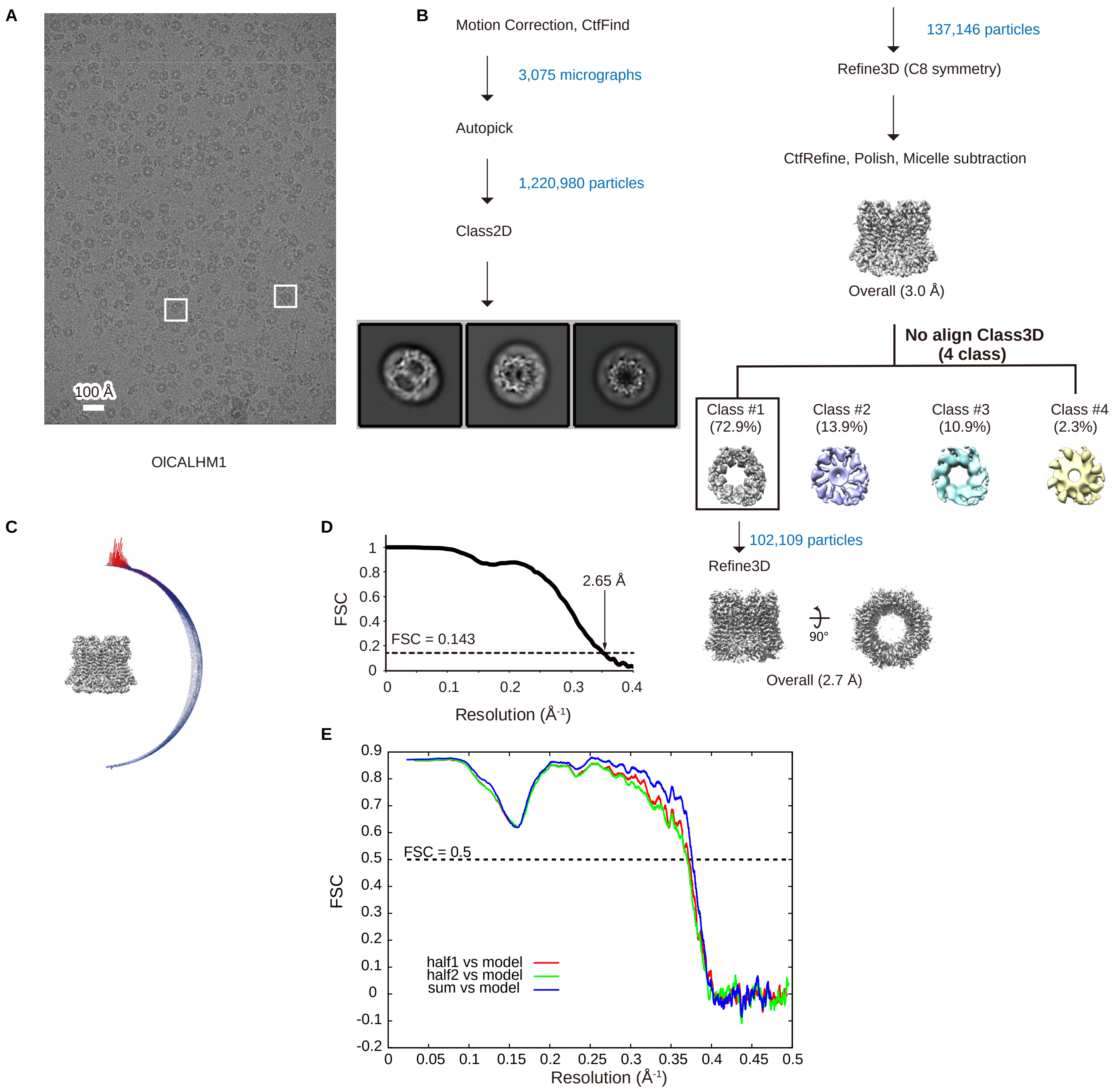

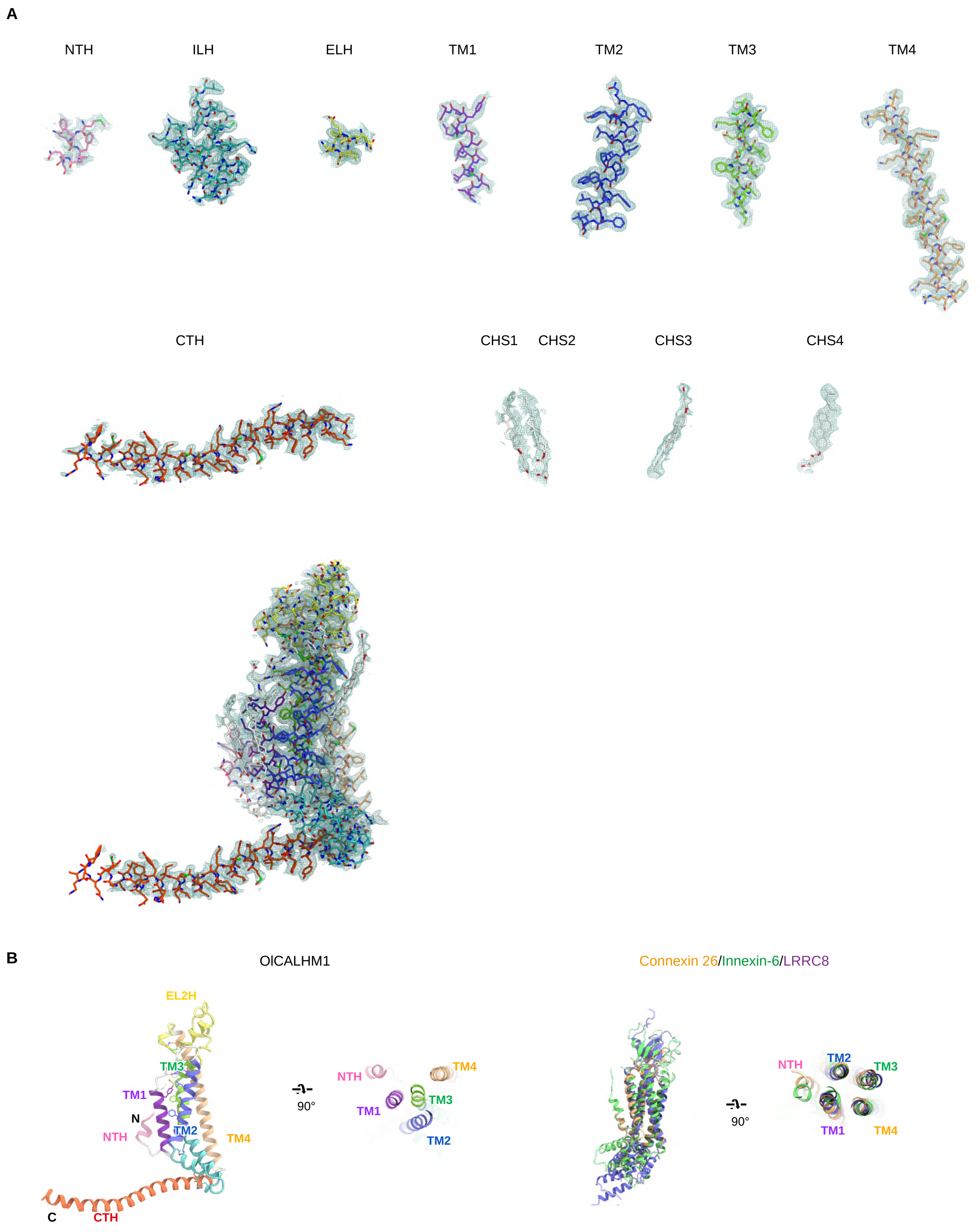

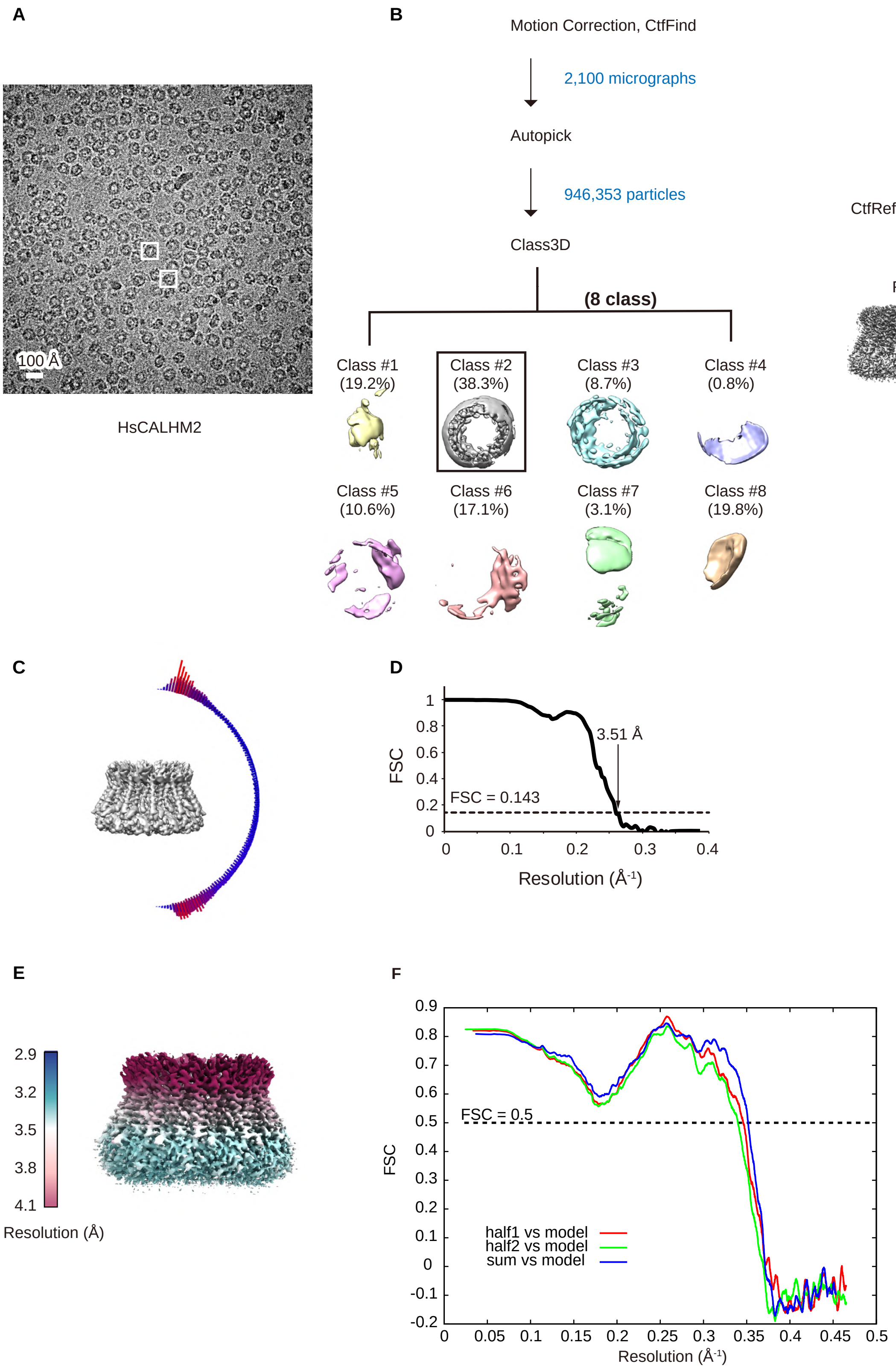

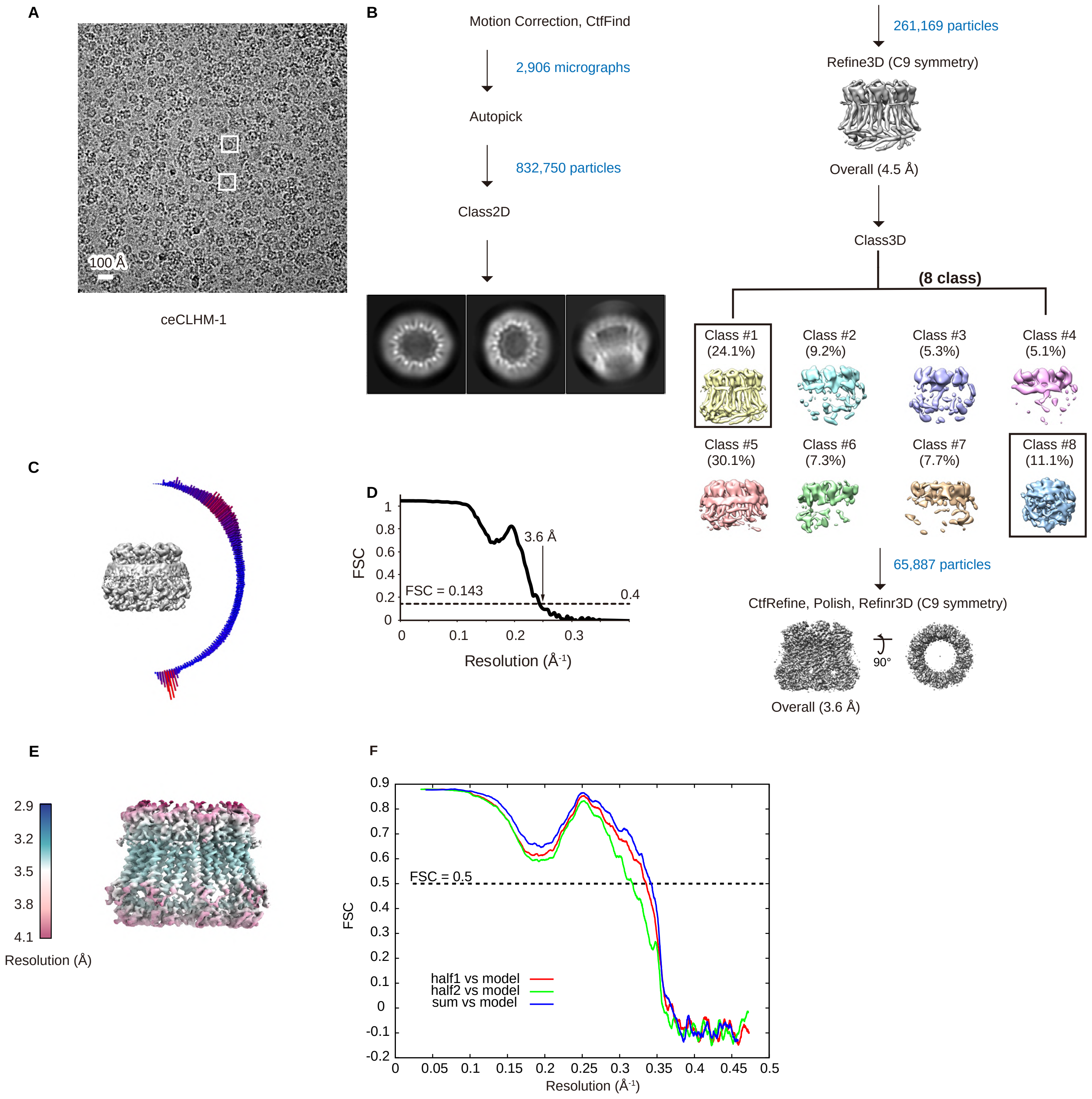

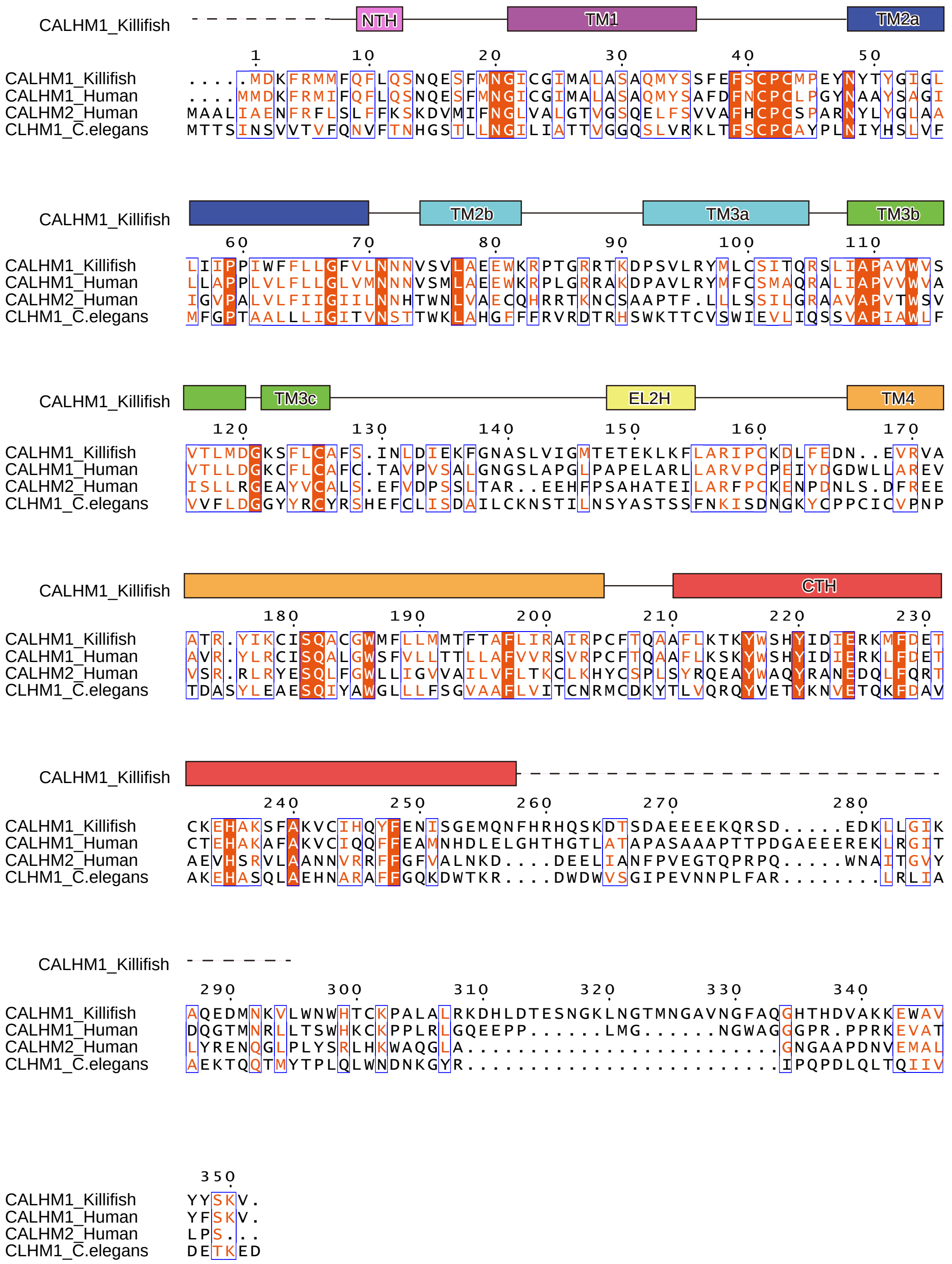

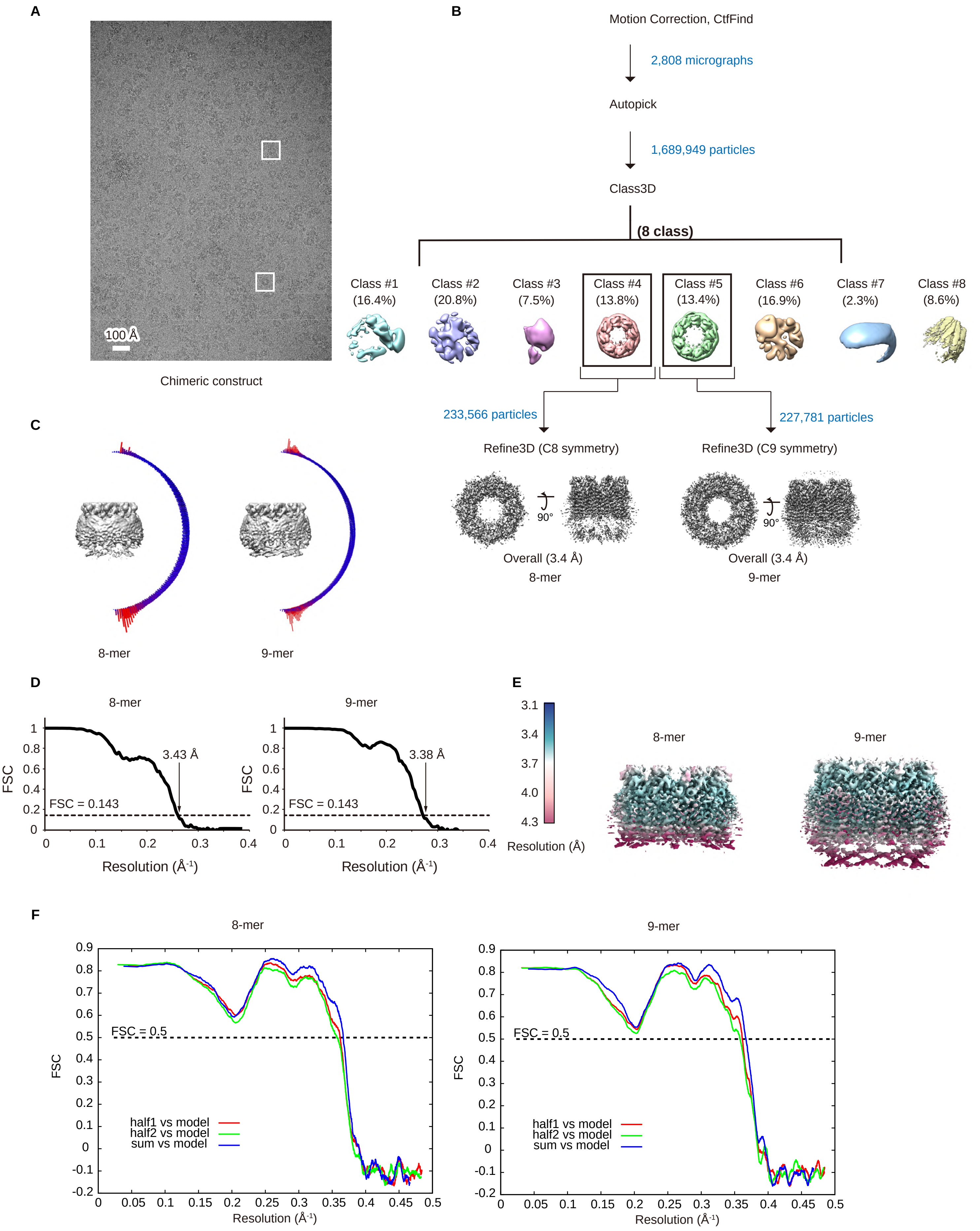

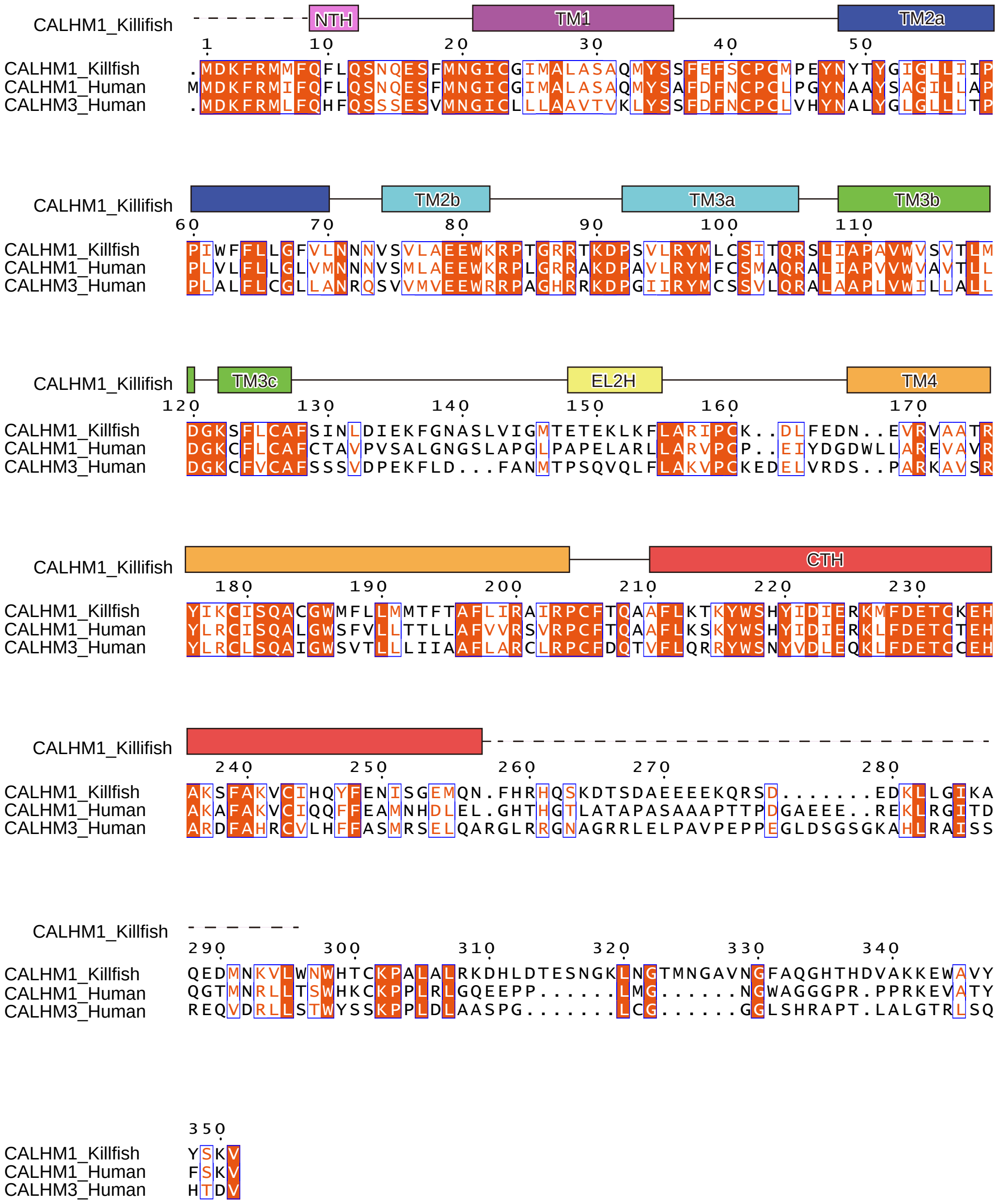
